# The Hao-Fountain syndrome gene USP7 restricts neurotropic orthoflavivirus entry through cell-intrinsic control of endosomal dynamics

**DOI:** 10.64898/2026.09.09.749922

**Authors:** Boris Bonaventure, Kyna Reyes, Marie-France Martin, Heng Pan, R Blake Richardson, Eva Bednarski, Alison W. Ashbrook, Allison Sowa, William Janssen, Oded Danzinger, Anastasija Cupic, Lisa Miorin, Charles M. Rice, Jean K. Lim, Brad R. Rosenberg, Matthew J. Evans, Jeffrey R. Johnson

## Abstract

Neurodevelopmental disorders are increasingly associated with immune phenotypes, including autoinflammation, immunodeficiency, and increased susceptibility to severe infection. To determine whether neurodevelopmental disorders-associated genes exert immune functions, we performed an arrayed siRNA screen targeting 28 genes with nonredundant cellular roles and assessed their effects on Zika virus (ZIKV) infection and innate immune pathways. We identified hits that intrinsically restrict ZIKV infection and modulate inflammatory pathways following infection. We further characterized the antiviral activity of the Hao-Fountain syndrome gene USP7, which potently restricts selected neurotropic orthoflaviviruses. USP7 inhibits ZIKV internalization before viral membrane fusion and genome release into the cytoplasm. Because USP7 plays a role in endosomal tubulation and recycling, we investigated whether endosomal recycling pathways restrict ZIKV infection. We identified the USP7-regulated E3 ubiquitin ligase TRIM27, as well as the recycling-associated Rab GTPases RAB11 and RAB35, as potent regulators of ZIKV infection. Infection assays using cell lines expressing pathogenic USP7 variants and primary fibroblasts from individuals with Hao-Fountain syndrome demonstrated that disease-associated USP7 mutations impair its antiviral activity and increase permissivity to ZIKV infection. These findings are consistent with recent case reports of severe viral infection during early life in individuals with Hao-Fountain syndrome. Collectively, our study identifies endosomal recycling pathways as important intrinsic restriction mechanisms against neurotropic orthoflaviviruses and nominates pathogenic USP7 variation as a candidate inborn error of immunity.

## INTRODUCTION

Host genetics strongly influence susceptibility to viral infectious diseases (*1*, *2*). The discovery of hundreds of inborn errors of immunity has shown that inherited or *de novo* mutations in immune genes can result in immunodeficiency, immune dysregulation, autoinflammatory disease, and life-threatening infections (*3–5*). The identification of these conditions has largely relied on clinical observation combined with genome or exome sequencing of affected individuals to pinpoint pathogenic variants in immune genes that explain their presentations (*6*, *7*). Over the past decade, these phenotype-to-genotype case studies have expanded the catalog of inborn errors of immunity to more than 500 genes, supporting the idea that additional pathogenic variants remain to be discovered (*3*).

Interestingly, recent studies examining the systemic consequences of inborn errors of immunity have highlighted links between immune genes defects and non-immune comorbidities (*8*, *9*). This suggests an unrecognized genetic overlap between genes at the frontline of immune defense and cellular pathways involved in diverse genetic diseases. To address this overlap, new genotype-to-phenotype approaches have been developed to probe the immune functions of genes that cause genetic disorders in which immune defects are not the primary clinical feature. For example, a recent study used a CRISPR-Cas9 screen in T cells to identify genes involved in immune cell differentiation among those implicated in inborn errors of metabolism, revealing a significant contribution of metabolic pathways to T-cell expansion (*10*).

Among described clinical overlaps, new observations have suggested connections between inborn errors of immunity and neurodevelopmental disorders (NDDs) (*11*, *12*). Indeed, it has been proposed that close to 20% of inborn errors of immunity also result in NDD symptoms such as microcephaly, epilepsy, autism spectrum disorders, and cognitive deficits. While the contribution of viral infection to the development of neurological symptoms is the subject of intense research and debate, the converse possibility, that defects in NDD genes could increase susceptibility to viral infection, remains largely unexplored (*13–15*). In this study, we aim to identify cell-intrinsic immune functions among genes involved in NDDs, to reveal new candidate inborn errors of immunity, and address the functional overlap between NDDs and immune disorders.

To do so, we performed an arrayed siRNA screen targeting 28 genes associated with nonredundant NDDs and cellular pathways relevant to viral replication. Using Zika virus (ZIKV) as a model orthoflavivirus, we identified NDD genes with cell-intrinsic antiviral and anti-inflammatory activities. Among these, the Hao-Fountain syndrome gene *USP7* emerged as a potent restriction factor for ZIKV and West Nile virus (WNV). Hao-Fountain syndrome is a multisystem neurodevelopmental disorder characterized by developmental delay, intellectual disability, autism spectrum disorder, speech impairment, and frequent gastrointestinal and sleep disturbances (*16*, *17*). Recent case reports also describe recurrent or unusually severe respiratory infections in affected individuals (*18–20*). Further characterization showed that USP7 restricts orthoflavivirus infection through its role with TRIM27 in endosomal recycling, providing first evidence that USP7 may be a candidate inborn error of immunity. Together, these findings uncover previously unrecognized immune functions of NDD-associated genes and identify endosomal recycling as a poorly defined cell-intrinsic antiviral defense pathway.

## RESULTS

### Arrayed siRNA screening identified antiviral genes

NDDs can be associated with immune dysregulation, including primary immunodeficiencies, autoinflammatory disease, and increased susceptibility to infections in some individuals (*11*, *21*, *22*). Conversely, inborn errors of immunity are frequently associated with neurodevelopmental conditions, supporting a functional overlap between immune pathways and genes involved in neurodevelopment (*12*). We hypothesized that certain genes involved in monogenic causes of NDDs could exert uncharacterized antiviral or immune functions. To test this hypothesis, we performed an arrayed siRNA screen targeting 28 NDD-associated genes to determine whether they exert antiviral activity against ZIKV or modulate innate immune signaling pathways. This gene set was curated from the Online Mendelian Inheritance in Man (OMIM) database, which catalogs genotype-phenotype associations for Mendelian genetic diseases (*23*). Selected genes are implicated in cellular pathways overlapping with the orthoflavivirus replication cycle, including endocytosis and endosomal trafficking, endoplasmic reticulum (ER) metabolism, Golgi function, and the ubiquitin-proteasome system (Fig. 1A). Our filtering strategy also prioritized ubiquitously expressed genes, avoided redundancy by selecting one gene per complex, and excluded genes previously known to impact orthoflavivirus infection or to have described immune functions (Fig. 1A). The gene set, their functions and associated disorders are listed in Table S1.

**Fig. 1.**
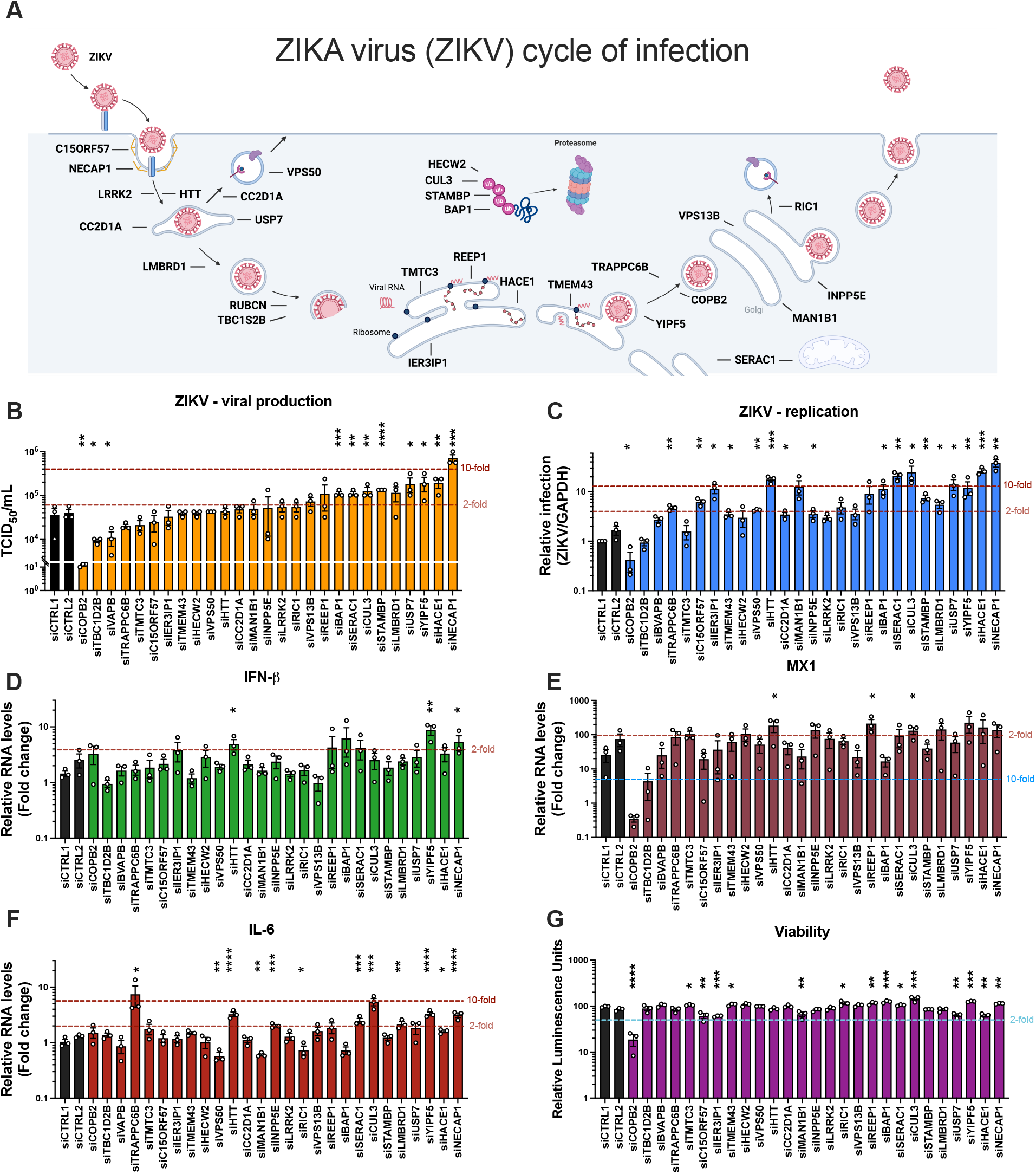
An arrayed siRNA screen identifies immune functions among NDD-associated genes. **(A)** Schematic of the Zika virus (ZIKV) replication cycle indicating candidate genes and their subcellular localization. **(B)** ZIKV production measured by TCID50 titration from supernatants, 24 h post-infection in siRNA-transfected control A549 cells (siCTRL1 and siCTRL2) or A549 cells depleted of the indicated target gene. **(C)** ZIKV infection efficiency measured by RT-qPCR (normalized to GAPDH) from cell lysates 24 h post-infection in siRNA-transfected A549 cells. **(D)** Relative expression of *IFN*-*β* measured by RT-qPCR (normalized to GAPDH and to non-infected siCTRL1 cells) from cell lysates 24 h post-infection in siRNA-transfected A549 cells. **(E)** Relative expression of *MX1* measured by RT-qPCR (normalized to GAPDH and to non-infected siCTRL1 cells) from cell lysates 24 h post-infection in siRNA-transfected A549 cells. **(F)** Relative expression of *IL-6* measured by RT-qPCR (normalized to GAPDH and to non-infected siCTRL1 cells) from cell lysates 24 h post-infection in siRNA-transfected A549 cells. **(G)** Cell viability expressed as relative luminescence units of luciferase measured using the Viral ToxGlow™ assay in siRNA-transfected A549 cells. Data information: (B–G) Mean ± SEM of three biological replicates. Multiple unpaired t-tests with false discovery rate (FDR) correction were performed to generate p-values. p-values are denoted as follows: *p < 0.05, **p < 0.01, ***p < 0.001, ****p < 0.0001.

A549 cells were transfected with siRNA pools targeting each gene, as well as two non-targeting siRNA pools used as negative controls. Transfected cells were then challenged with the ZIKV strain MR-766, and infection phenotypes, along with innate immune and inflammatory responses, were assessed 24 h post-infection (Fig. 1, B to F). Viral production was quantified by TCID50 titration on Vero cells and viral RNA replication by RT-qPCR (Fig. 1,B and C). Silencing of 8 targets: *NECAP1*, *HACE1*, *YIPF5*, *USP7*, *STAMBP*, *CUL3*, *SERAC1* and *BAP* significantly increased ZIKV production by at least 2-fold. Depletion of 6 of those genes: *NECAP1*, *HACE1*, *YIPF5*, *USP7*, *CUL3* and *SERAC1* also led to at least a 10-fold increase in viral replication, suggesting that they exert potent antiviral activity (Fig. 1C). NECAP1 is a clathrin-accessory protein that regulates endocytosis and is implicated in early infantile epileptic encephalopathy (*24*, *25*). HACE1 is a HECT-type E3 ubiquitin ligase that catalyzes ubiquitin transfer to substrates, leading to their degradation or functional modulation; pathogenic mutations in *HACE1* cause hereditary spastic paraplegia, a degenerative disorder characterized by muscle weakness in the legs (*26*, *27*). YIPF5 is an ER-membrane protein that orchestrates ER-to-Golgi trafficking and regulates SURF4-mediated ER export; pathogenic mutations cause microcephaly associated with epilepsy and diabetes syndrome (*28*, *29*). USP7 is a ubiquitin hydrolase that removes ubiquitin tags from substrates and is mutated in the neurodevelopmental disorder known as Hao-Fountain syndrome (*16*, *17*, *30*). CUL3 is a scaffold protein for Cullin-RING E3 ubiquitin ligase complexes and is implicated in a neurodevelopmental disorder associated with autism spectrum disorder (*31*, *32*). SERAC1 regulates intracellular cholesterol trafficking, and pathogenic mutations lead to encephalopathy and mitochondrial disorders (*33*, *34*).

Silencing of 11 additional genes, including *LMBRD1*, *STAMP*, *BAP1*, *REEP1*, *NPP5E*, *CC2D1A*, *HTT*, *TMEM43*, *IER3IP1*, *C15ORF73* and *TRAPPC6B* also significantly promoted ZIKV RNA replication but with a modest effect or no effect on viral production (Fig. 1B and C). The innate immune response of infected cells was evaluated by measuring transcription of *interferon* (*IFN)β* and *MX1*, a prototypic interferon-stimulated gene (ISG) (Fig. 1D and E). Interestingly, genes exhibiting antiviral activity did not significantly altered *IFNβ* or *MX1* expression upon infection suggesting these antiviral effects are independent on the type-I IFN response. *HTT* silencing significantly enhanced both *IFNβ* and *MX1* transcription, suggesting a potential role in innate immune sensing. Depletion of *NECAP1* and *YIPF5* significantly enhanced transcription of *IFNβ* but not *MX1*, which suggests a potential function in the innate sensing. Conversely, depletion of *BAP1* and *CUL3* significantly upregulated only *MX1* transcription. Because depletion of these 5 genes significantly promoted ZIKV infection, their apparent effects on the type-I IFN pathway might instead reflect increased abundance of viral nucleic acids available for sensing.

Several gene knockdowns had a significant impact on *IL-6* transcriptional induction upon ZIKV infection (Fig. 1F). Silencing of *NECAP1*, *HACE1*, *YIPF5*, *LMBRD1*, *CUL3*, *RIC1*, *SERAC1*, *NPP5E*, *HTT* and *TRAPPC6B* potentiated *IL-6* transcription, suggesting regulatory roles in inflammatory responses. TRAPPC6B and CUL3 had the most notable anti-inflammatory effect, as their depletion increased *IL-6* transcription by 7-fold and 5-fold, respectively (Fig. 1F). Conversely, depleting *BAP1*, *RIC1*, *LRRK2*, *MAN1B1*, and *VPS50* modestly but significantly decreased *IL-6* transcription, indicating potential pro-inflammatory functions of these genes. Cell viability assays confirmed little to no cytotoxicity associated with the siRNAs, except for those targeting *COPB2* (Fig. 1G), which likely explains why COPB2-depleted cells were significantly less infected by ZIKV (Fig. 1, A and B). Altogether, this targeted siRNA screen identified several NDD-associated genes with intrinsic antiviral activity against ZIKV and provided candidate genes that may be involved in virus-induced inflammation.

### USP7 selectively regulates orthoflavivirus infection in an intrinsic manner

Based on our screen, we prioritized USP7 because Hao-Fountain syndrome, caused by heterozygous pathogenic USP7 variants, has been reported in a larger patient cohort than disorders linked to other candidate genes (*17*). USP7 is known as a herpesvirus cofactor that stabilizes viral proteins (*35–37*). It is notably hijacked by the HSV E3 ubiquitin ligase ICP0 to prevent its degradation and thereby support productive viral infection (*38*, *39*). However, to our knowledge, the antiviral functions of USP7 and its role in orthoflavivirus infection remain undefined. We first validated its antiviral activity against ZIKV and found that siRNA-mediated depletion of USP7 in A549 cells potently enhanced infection with the ZIKV strain MR-766 (Fig. 2A). USP7-depleted cells infected with increasing multiplicity of infection (MOI) of ZIKV for 24 h showed a robust increase in infection as measured by flow cytometry with intracellular staining of the envelope (E) protein. Viral infection, quantified by RT-qPCR and by TCID50 titration, revealed a 5-to 10-fold increase in viral RNA levels and viral production, respectively (Fig. 2A). Of note, USP7 silencing was approximately 90% effective and did not affect cell viability (Fig. 2, B and C and fig. S1A). We next asked whether the antiviral activity of USP7 was specific to ZIKV or extended to other orthoflaviviruses. USP7-depleted A549 cells were challenged with other mosquito-borne orthoflaviviruses, and we found that USP7 exerted an antiviral effect on infection with WNV strain NY99 (Fig. 2D). Viral infection, measured by RT-qPCR and TCID50 24 h post-infection, showed an approximately 10-fold increase in viral RNA levels and WNV production. Consistently, USP7 CRISPR-Cas9 knockout significantly increased WNV infection despite only modest knockout efficiency at the population level (fig S1, B and C). Interestingly, USP7 depletion did not impact infection by dengue virus serotype 2 (DENV-2) strain 16681 or yellow fever virus (YFV) strain 17D, as assessed by intracellular staining, RT-qPCR, and TCID50 titration (Fig. 2, E and F). These findings suggest that USP7 exerts antiviral activity selectively against specific neurotropic flaviviruses. Then, we validated the antiviral activity of USP7 in another cell model, hepatocyte-derived Huh7.5 cells, where USP7 depletion using individual siRNAs produced a similar antiviral phenotype proportional to the silencing efficiency (Fig. 2, G and H).

**Fig. 2.**
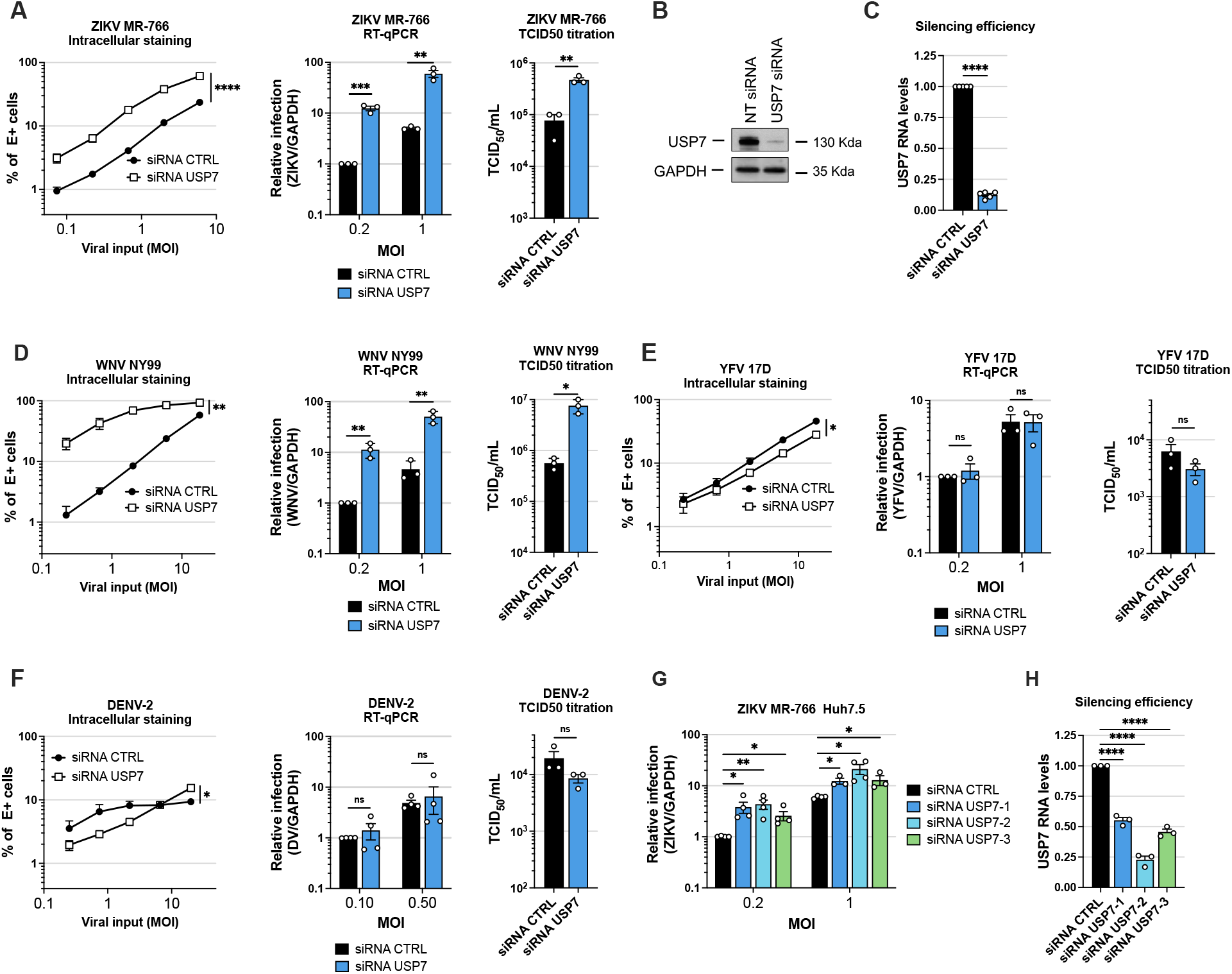
USP7 depletion promotes ZIKV and WNV infection in cell lines. **(A)** Zika virus (ZIKV) strain MR-766 infection rates at 24 h post infection were quantified in A549 cells transfected with non-targeting siRNA pools (siRNA CTRL) or siRNA pools targeting USP7 (siRNA USP7). Infection was measured by intracellular staining of the viral Envelop (E) protein and flow cytometry analysis at different multiplicities of infection (MOI), by RT-qPCR at two MOI, and by TCID50 titration of supernatants from cells infected at an MOI of 0.2. Simple linear regression analysis on log-transformed data and multiple unpaired t-tests with false discovery rate correction. **(B)** USP7 silencing efficiency in A549 cells compared to non-targeting siRNA pools (NT siRNA) measured by immunoblot. **(C)** USP7 silencing efficiency in A549 cells measured by RT-qPCR. Unpaired t-test. **(D)** West Nile virus (WNV) strain NY99 infection rates at 24 h post infection were quantified in A549 cells transfected with siRNA CTRL or siRNA USP7. Infection was measured by intracellular staining of the viral E protein and flow cytometry analysis at different MOI, by RT-qPCR at two MOIs, and by TCID50 titration of supernatants from cells infected at an MOI of 0.2. Simple linear regression analysis on log-transformed data and multiple unpaired t-tests with false discovery rate correction. **(E)** Yellow fever virus (YFV) strain 17D infection rates at 24 h post infection were quantified in A549 cells transfected with siRNA CTRL or siRNA USP7. Infection was measured by intracellular staining of the viral E protein and flow cytometry analysis at different MOI, by RT-qPCR at two MOI, and by TCID50 titration of supernatants from cells infected at an MOI of 0.2. Simple linear regression analysis on log-transformed data and multiple unpaired t-tests with false discovery rate correction. **(F)** Dengue-2 virus (DENV-2) strain 16681 infection rates at 48 h post infection were quantified in A549 cells transfected with siRNA CTRL or siRNA USP7. Infection was measured by intracellular staining of the viral E protein and flow cytometry analysis at different MOIs, by RT-qPCR at two MOI, and by TCID50 titration of supernatants from cells infected at an MOI of 0.5. Simple linear regression analysis on log-transformed data and multiple unpaired t-tests with false discovery rate correction. **(G)** ZIKV strain MR-766 infection rates at 24 h post infection were quantified in Huh7.5 cells transfected with control siRNA (siRNA CTRL) or individual siRNAs targeting USP7 (siRNA USP7-1, -2, -3). Infection was assessed by RT-qPCR at two MOI. Multiple unpaired t-tests with false discovery rate correction. **(H)** USP7 silencing efficiency in Huh7.5 cells measured by RT-qPCR. Unpaired t-test. Data information: (A, C-H) Mean ± SEM of three or four biological replicates. p-values are denoted as follows: ns, not significant, *p < 0.05, **p < 0.01, ***p < 0.001, ****p < 0.0001.

We next asked whether this antiviral effect of USP7 was mediated through the IFN response, as impaired innate immune activation can lead to higher viral replication. Upon ZIKV and WNV infection, USP7 depletion did not reduce transcription of prototype ISGs such as *MX1* or *ISG15* but instead increased their expression (fig. S1, D and E). We interpret this enhanced ISG transcription to be a consequence of higher viral replication that favors pathogen recognition. To rule out a contribution of antiviral ISGs to this phenotype, we pre-treated cells with a combination of TBK1 and pan-JAK inhibitors to abolish IFN production and downstream IFN signaling during infection. *USP7* silencing still greatly promoted WNV infection in inhibitor-treated cells, indicating that its antiviral function is IFN-independent (fig S1F). Efficient TBK1/JAK inhibition was confirmed by RT-qPCR, as *MX1* and *ISG15* mRNA levels in infected samples were comparable to those in non-infected cells (fig S1, G and H).

Altogether, these data suggest that USP7 selectively inhibits neurotropic mosquito-borne orthoflaviviruses in a type-I IFN-independent manner.

### USP7 regulates orthoflavivirus internalization before viral fusion

Next, we sought to investigate at which step of the viral life cycle USP7 inhibits orthoflavivirus infection. To determine whether USP7 affects viral propagation or inhibits infection during the first replication cycle, A549 cells were infected with ZIKV for four hours, and the infection medium was supplemented with ammonium chloride (NH_4_Cl), which blocks infection by newly produced virions (Fig. 3A). The antiviral effect exerted by USP7 was not impacted by NH_4_Cl treatment post-infection, as shown by quantification of ZIKV infection by RT-qPCR and TCID50 titration (Fig.3A). This indicates that USP7 impacts ZIKV infection during the first round of replication and does not affect viral propagation at the 24 h post-infection time point. We determined whether USP7 depletion impacted virus binding by performing a binding assay in which cells and viral inoculum were incubated at 4 °C for one hour before extensive washing with PBS and lysis for RT-qPCR analysis of viral RNA (Fig. 3B). USP7 depletion did not affect virus binding to the cells. However, internalization assays, in which cells were incubated at 37 °C for short time points after virus binding, showed that virus internalization was greatly enhanced in the absence of USP7 as early as one hour post-infection (Fig. 3C). This suggests that USP7 blocks ZIKV internalization into cells without affecting viral binding. To confirm that we were quantifying incoming viral particles that had not undergone replication, internalization assays were conducted in cells pre-treated with NH_4_Cl, bafilomycin, or cycloheximide to inhibit envelope fusion, endosomal acidification, or translation, respectively, and then infected with ZIKV for 4 h (Fig. 3D). The USP7 depletion associated phenotype was not affected by any of these treatments, confirming that our internalization assay primarily quantifies incoming viral genomes and suggesting that USP7 acts upstream of viral envelope fusion and release of the viral genome into the cytoplasm. To determine whether, in the absence of USP7, virions entered via similar routes as in control conditions, we pre-treated cells with Pitstop2, an inhibitor of clathrin-mediated endocytosis, and infected them with ZIKV for 24 h (Fig S2A) (*40*). Upon USP7 depletion, ZIKV infection was still inhibited by Pitstop2 pre-treatment, indicating that virions do not bypass clathrin-mediated endocytosis when USP7 is depleted. Altogether, these experiments show that USP7 exerts an intrinsic pressure early in the infection life cycle, by limiting virus internalization without modifying the entry route of orthoflaviviruses.

**Fig. 3.**
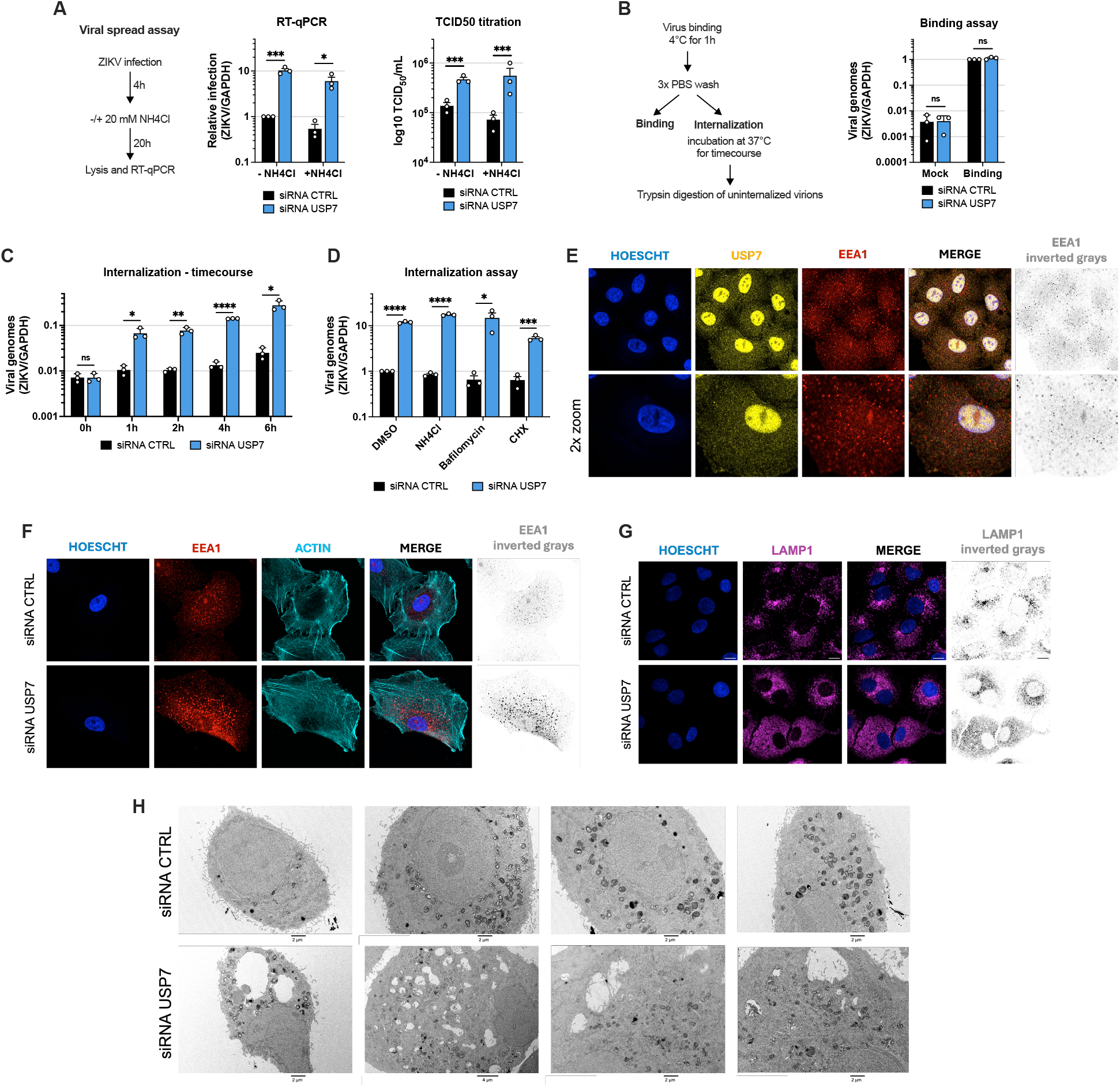
USP7 limits ZIKV internalization and regulates early and late endosome organization. **(A)** Zika virus (ZIKV) strain MR-766 infection at 24 h post-infection in A549 cells transfected with non-targeting siRNA pools (siRNA CTRL) or siRNA pools targeting USP7 (siRNA USP7), with or without post-treatment with ammonium chloride (NH4Cl) to block additional rounds of infection. Infection was measured by RT-qPCR at indicated multiplicity of infection (MOI), and by TCID50 titration of supernatants from cells infected at an MOI of 0.2. Statistical analysis was performed using multiple unpaired t-tests with false discovery rate correction. **(B)** For binding assays, siRNA CTRL or siRNA USP7 A549 cells were incubated with ZIKV at an MOI of 10 at 4 °C for 1 h. After extensive washing, cell-bound virus was quantified by RT-qPCR. Statistical analysis was performed using multiple unpaired t-tests with false discovery rate correction. **(C)** For internalization assays, ZIKV (MOI 10) was first bound to siRNA CTRL or siRNA USP7 A549 cells at 4 °C for 1 h and then shifted to 37 °C to trigger viral entry. At the indicated time points, cells were trypsinized to remove surface-bound virus, lysed, and total RNA extracted to quantify internalized virus by RT-qPCR. Viral RNA levels were normalized to the relative amount of bound virus, and RNA levels at the 0 h time point were similar to those of the mock condition shown in Figure 3B. Statistical analysis was performed using multiple unpaired t-tests with false discovery rate correction. **(D)** Internalization assays were performed in siRNA CTRL and siRNA USP7 A549 cells pre-treated with NH4CL, Bafilomycin, Cycloheximide (CHX), or equivalent volume of DMSO as negative control. Statistical analysis was performed using multiple unpaired t-tests with false discovery rate correction. **(E)** Confocal microscopy of A549 cells stained with anti-EEA1 (red and inverted gray) and anti-USP7 (yellow) antibodies, and Hoescht for nuclei (blue). Representative images from two independent experiments. Top: scale bar: 10 μm. Bottom: scale bar: 20 μm. **(F)** Confocal microscopy of siRNA CTRL and siRNA USP7 A549 cells stained with anti-EEA1 (red and invested gray), phalloidin for F-actin (cyan), and Hoescht for nuclei (blue). Representative images from three independent experiments Scale bar: 10 µm. **(G)** Confocal microscopy of siRNA CTRL and siRNA USP7 A549 cells stained with anti-LAMP1 (purple) and Hoescht for nuclei (blue). Representative images from two independent experiments Scale bar: 10 µm. **(H)** Electron micrographs of siRNA CTRL and siRNA USP7 A549 cells. Scale bars: 2 or 4 µm, as indicated in the images. Data information: **(A-D)** Mean ± SEM of three biological replicates. p-values are denoted as follows: ns, not significant, *p < 0.05, **p < 0.01, ***p < 0.001, ****p < 0.0001.

Endogenous USP7 localization analyzed by confocal microscopy showed that USP7 is present in both the nucleus and cytoplasm (Fig. 3E). Because USP7 inhibits ZIKV entry into cells, we next examined whether USP7 colocalizes with endosomal compartments involved in orthoflavivirus trafficking, such as early and late endosomes. USP7 did not colocalize with markers of early endosomes (EEA1) or late endosomes/lysosomes (LAMP1), suggesting that USP7 does not directly associate with endosomal compartments involved in orthoflavivirus uptake and trafficking during viral entry (Fig. 3E and fig. S2B). However, USP7 depletion in A549 cells had marked effects on the morphology of endosomes. USP7 depletion increased the number and intensity of early endosomal vesicles, as shown by EEA1 staining, and profoundly disrupted the polarity of the late endosomal compartment (Fig. 3, F and G). Electron microscopy revealed striking subcellular morphological changes in USP7-depleted cells, including the appearance of large, low-density vesicles (Fig. 3H). USP7-depleted cells also showed a potential accumulation of electron-dense structures, which we interpret as likely lysosomes (fig. S2C). Overall, this microscopy analysis indicates that USP7 depletion affects endosomal compartment organization and triggers the enlargement of subcellular structures that we hypothesize to be enlarged vesicles.

### TRIM27 regulates orthoflavivirus infection and is stabilized by USP7

USP7 has been shown to stabilize and regulate the activity of the MAGEL2-USP7-TRIM27 (MUST) endosomal complex, which tunes WASH ubiquitination (*16*). In this complex, TRIM27 is a well-characterized USP7 substrate that is degraded by the proteasome upon USP7 depletion. The MUST complex associates with the retromer and promotes WASH-dependent actin polymerization on early endosomes, thereby elongating endosomal tubules and supporting vesicle scission (*41*). The retromer complex recycles endosomal cargo, primarily via endosome-to-Golgi retrograde transport, but also through endosome-to-plasma-membrane recycling pathways (*42*). Because USP7 affects orthoflavivirus internalization and endosomal sorting machineries regulate endosomal trafficking, we next asked whether TRIM27 and the retromer complex exert antiviral activity against USP7-sensitive flaviviruses.

A549 cells were transfected with siRNA pools targeting USP7, TRIM27, or VPS35, the central core retromer subunit, to determine whether these complexes regulate orthoflavivirus infection. TRIM27 depletion promoted WNV infection to a similar extent as USP7 silencing, whereas VPS35 depletion did not affect viral infection (Fig. 4A and fig S3A). According to gene expression databases, A549 cells express low or undetectable levels of MAGEL2, which could explain a retromer-independent function of USP7 and TRIM27 (*43*). These results suggest that USP7 and TRIM27 may act together to limit orthoflavivirus infection independently of the retromer. TRIM27 auto-ubiquitination leads to its destabilization, and previous studies have shown that USP7 stabilizes TRIM27 by preventing its degradation (*16*). Consistent with this, we found that TRIM27 expression was regulated by USP7, whereas VPS35 depletion was not (Fig. 4B). As observed for USP7, TRIM27 localized to both the nucleus and the cytoplasm in punctate structures (fig S3B). In A549 cells ectopically expressing FLAG-TRIM27, USP7 and TRIM27 co-localized in the nucleus as well as in cytoplasmic granules (Fig. 4C).

**Fig 4.**
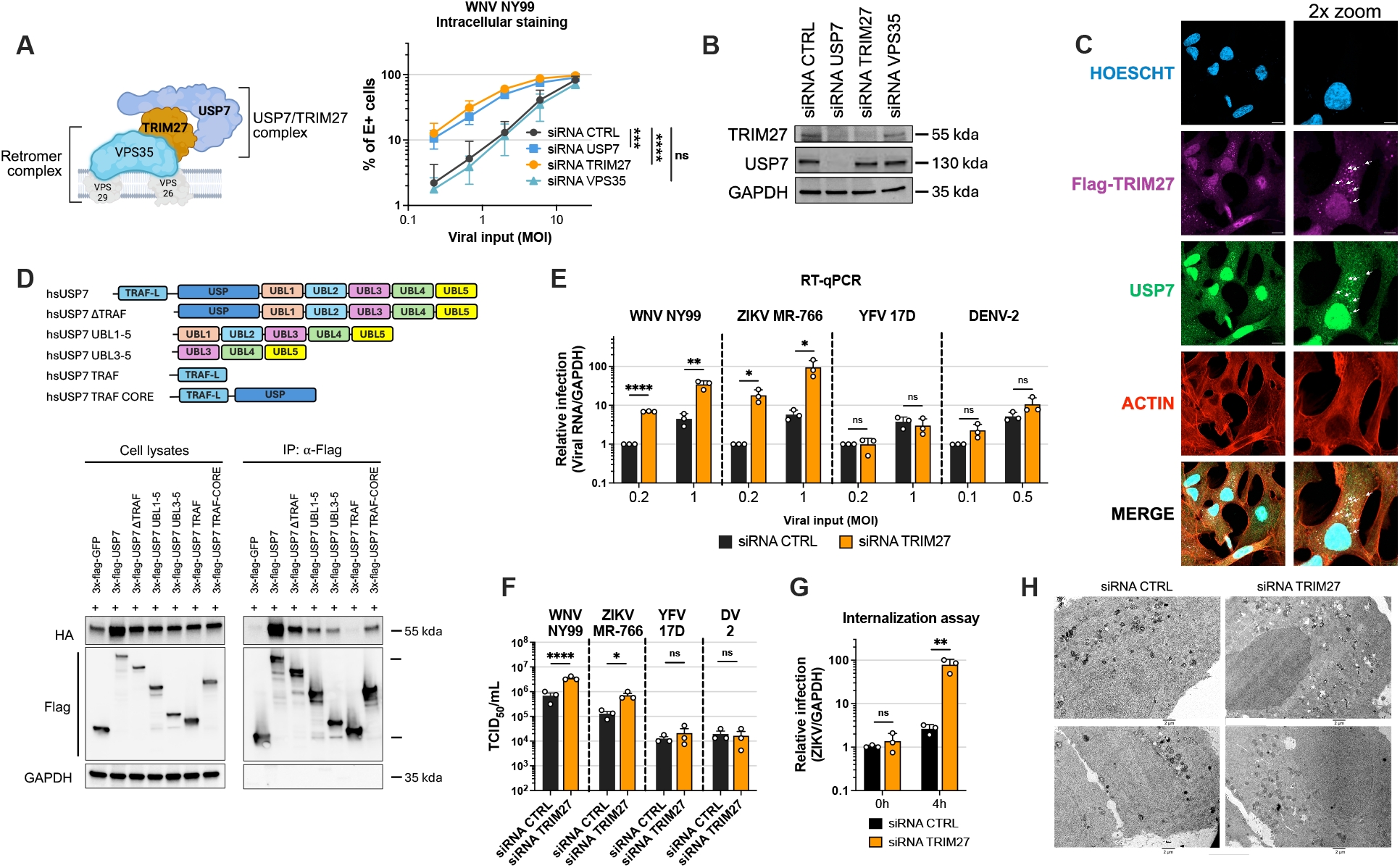
USP7 stabilizes and interacts with TRIM27, a ubiquitin E3 ligase with antiviral activity.(A) Schematic depicting the USP7/TRIM27 complex interacting with the retromer complex composed of the VPS35, VPS29, and VPS26 subunits. West Nile virus (WNV) strain NY99 infection was measured at 24 h post-infection in A549 cells transfected with non-targeting siRNA pools (siRNA CTRL) or siRNA pools targeting USP7 (siRNA USP7), TRIM27 (siRNA TRIM27), or VPS35 (siRNA VPS35). Infection was measured by intracellular staining of the viral Envelop (E) protein and flow cytometry analysis at different MOI. Statistical analysis was performed using simple linear regression analysis. **(B)** USP7 and TRIM27 silencing efficiency in siRNA transfected A549 cells measured by immunoblot. **(C)** Confocal microscopy of A549 cells expressing Flag-TRIM27 stained with anti-flag (purple), anti-USP7 (green) antibodies, phalloidin for F-actin (red), and Hoescht for nuclei. Left: scale bar: 10 µm. Zoom right: scale bar: 20 µm. **(D)** Top: Schematic representation of USP7 domain architecture and truncation mutants. The N-terminal region of USP7 contains a TRAF-like domain (TRAF-L), followed by the catalytic ubiquitin-specific protease domain (USP) and five consecutive ubiquitin-like domains (UBL1–5). Bottom: Co-immunoprecipitation experiments in HEK293T cells transfected to express 3×FLAG-GFP (negative control), 3×FLAG-USP7, or the indicated USP7 truncation mutants together with HA-TRIM27. At 24 h post-transfection, cells were lysed, and FLAG-tagged proteins were immunoprecipitated using anti-FLAG magnetic beads, and co-immunoprecipitated proteins were detected by immunoblot. Representative immunoblot from two independent experiments. **(E)** WNV NY99, Zika virus (ZIKV) MR-766, Yellow fever virus (YFV) 17D and Dengue virus 2 (DENV-2) 16681 infection rates were quantified in A549 cells transfected with siRNA CTRL or siRNA TRIM27 at 24 h post infection. Cells were infected at the indicated multiplicity of infection (MOI), and infection was measured by RT-qPCR. Multiple unpaired t-tests with false discovery rate correction. **(F)** WNV NY99, ZIKV MR-766, YFV 17D, and DENV-2 16681 viral productions were quantified from A549 cells transfected with siRNA CTRL or siRNA TRIM27 at 24 h post-infection. Cells were infected at an MOI of 0.2, and infection was measured by TCID50 titration of cell culture supernatants. Multiple unpaired t-tests with false discovery rate correction. **(G)** Internalization assays were performed by incubating siRNA CTRL or siRNA TRIM27 A549 cells with ZIKV at an MOI of 10, and by stopping internalization after 0 or 4 h. Statistical analysis was performed using unpaired t-tests with false discovery rate correction. **(H)** Electron micrographs of siRNA CTRL and siRNA TRIM27 A549 cells. Scale bars: 2 µm. Data information: (A, E-G) Mean ± SEM of three biological replicates. p-values are denoted as follows: ns, not significant, *p < 0.05, **p < 0.01, ***p < 0.001, ****p < 0.0001.

Next, we sought to confirm the physical interaction between USP7 and TRIM27 and to determine which USP7 domains mediate this interaction. Co-transfection of FLAG-tagged USP7 and HA-tagged TRIM27 in HEK293T cells, followed by USP7 immunoprecipitation, showed that TRIM27 physically interacts with USP7 (Fig. 4D). USP deubiquitinases rely on a catalytic core flanked by additional domains that confer substrate and functional specificity. USP7 is structurally distinct among USPs: its N-terminus contains a TRAF-like domain connected to the USP core, and its C-terminus comprises five successive ubiquitin-like (UBL) domains. Co-transfection of FLAG-tagged USP7 truncation mutants (Fig. 4C) with HA-tagged TRIM27 and immunoprecipitation of FLAG-USP7 demonstrated that TRIM27 binds the USP7 core domain as well as the UBL domains but does not bind the TRAF-like domain. This binding interface is consistent with previous in vitro USP7-TRIM27 structure-function studies and with USP7 enzymatic characterization showing that substrate binding triggers a conformational change that connects the UBL4-5 domains with the catalytic core to stabilize substrate interactions (*16*, *44*).

To determine whether TRIM27 exhibits the same antiviral specificity toward orthoflaviviruses as USP7, we infected TRIM27-depleted A549 cells with ZIKV MR-766, WNV NY99, YFV 17D, and DENV-2 16681 (Fig. 4E). Similar to USP7 depletion, TRIM27 knockdown increased WNV and ZIKV replication by more than 10-fold, with a proportional increase in viral production but did not significantly affect YFV or DENV-2 infection (Fig. 4, E and F). This TRIM27-mediated antiviral activity was also type I IFN-independent, as pre-treatment with pan-JAK and TBK1 inhibitors did not alter this phenotype (Fig. S3C). We investigated if TRIM27 inhibited orthoflavivirus internalization and showed that TRIM27 depletion promoted ZIKV entry by 50-fold (Fig. 4G). Overall, these data highlight TRIM27’s potent antiviral activity and reveal that the USP7-TRIM27 complex intrinsically restricts orthoflavivirus entry, independently of retromer function.

To determine whether TRIM27 depletion affects endosomal compartment structure, we stained TRIM27-depleted cells with EEA1 and LAMP1 as markers of early and late endosomes, respectively (fig S3, D and E). Similar to USP7 depletion, TRIM27 silencing led to an increased number and intensity of early endosomes and a disrupted organization of the LAMP1-positive compartment. However, the effects of TRIM27 depletion on early and late endosome morphology and organization were less pronounced than those observed in USP7-depleted cells (Fig. S3, D and E). Electron micrographs of TRIM27-depleted cells showed a milder, yet similar, phenotype to USP7-depleted cells, characterized by an accumulation of large, low-density vesicles (Fig. 4H). Altogether, these data indicate that USP7 controls the stability of TRIM27, and that depletion of either protein alters early and late endosomal compartments by affecting their morphology and organization.

### Selected endosomal recycling pathways restrict ZIKV infection

Because the USP7-TRIM27 complex participates in endosomal recycling and its depletion perturbs endosomal compartmentalization, we asked whether specific endosomal recycling pathways restrict infection by USP7-sensitive orthoflaviviruses. We previously showed that depletion of VPS35, a core retromer subunit, does not affect WNV infection (Fig. 4A), suggesting that retromer-dependent recycling is not required for this restriction. We therefore hypothesized that alternative recycling routes might limit orthoflavivirus infection in a manner similar to USP7 and TRIM27. To test this, we depleted A549 cells of key endosomal recycling regulators, including sorting nexin 17 (SNX17), which functions in the SNX17-Retriever pathway that drives endosome-to-plasma-membrane recycling, and the Rab GTPases Rab4, Rab11, and Rab35 (*45*– *47*). Rab4 controls fast recycling from early/sorting endosomes to the plasma membrane, Rab11 regulates recycling through perinuclear recycling endosomes and contributes to constitutive exocytosis, and Rab35 operates at the plasma membrane and endosomes to coordinate fast recycling and exosome release (*47*). Silencing Rab4A, Rab11A, and Rab35A in A549 cells significantly enhanced ZIKV infection, with Rab11A and Rab35A depletion reaching levels comparable to TRIM27 knockdown (Fig. 5A), indicating that intact Rab-dependent recycling pathways act as intrinsic barriers to ZIKV replication. In contrast, SNX17 knockdown had no effect on ZIKV, demonstrating that recycling pathways are not uniformly antiviral for this virus. YFV 17D, which is insensitive to USP7 or TRIM27 depletion, was likewise not enhanced by Rab4A, Rab11A, or Rab35A knockdown and was even reduced upon Rab35A depletion, consistent with studies reporting a partial dependence of YFV on endosomal recycling (*48*) (Fig. 5B). Conversely, SNX17 depletion significantly increased YFV 17D infection, although not as strongly as the effect of TRIM27 on ZIKV infection. Silencing efficiencies were validated by RT-qPCR and ranged from 70% to 95% knockdown (fig S4A). Together, these data indicate that ZIKV is markedly restricted by specific Rab-dependent recycling pathways, whereas YFV 17D displays an opposite sensitivity profile, suggesting that orthoflaviviruses differ in their interactions with recycling endosomal routes and may exploit more diverse entry and trafficking pathways than previously appreciated.

**Figure 5.**
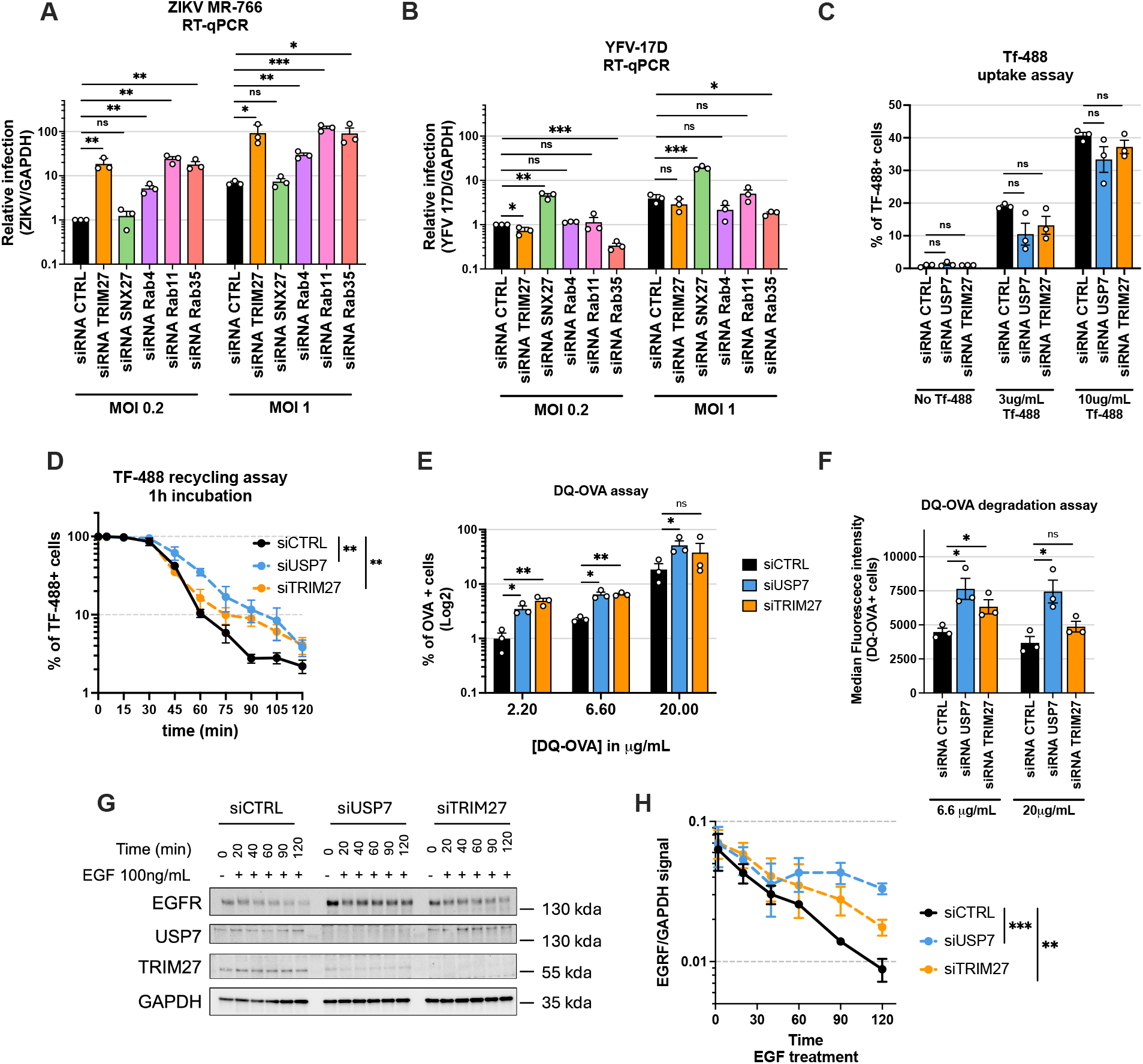
Defects in endosomal recycling factors promote ZIKV infection and dysregulate endosomal trafficking. **(A)** Zika virus (ZIKV) strain MR-766 infection at 24 h post-infection in A549 cells transfected with non-targeting siRNA pools (siRNA CTRL) or siRNA pools targeting USP7 (siRNA USP7), TRIM27 (siRNA TRIM27), SNX27 (siRNA SNX27), Rab4 (siRNA Rab4), Rab11 (siRNA Rab11) or Rab35 (siRNA Rab35). Infection was measured by RT-qPCR at indicated multiplicity of infection (MOI). Statistical analysis was performed using multiple unpaired t-tests with false discovery rate correction. **(B)** Yellow fever virus (YFV) strain 17D infection at 24 h post-infection in siRNA CTRL, siRNA USP7, siRNA TRIM27, siRNA SNX27, siRNA Rab4, siRNA Rab11 or siRNA Rab35 A549 cells. Infection was measured by RT-qPCR at indicated MOI. Statistical analysis was performed using multiple unpaired t-tests with false discovery rate correction. **(C)** Transferrin (Tf)-488 uptake assays in siRNA CTRL, siRNA USP7, or siRNA TRIM27 A549 cells. Cells were incubated with the indicated concentrations of Tf-488 for 30 min at 4 °C and then shifted for 10 min to 37 °C to allow Tf-488 internalization. The percentage of Tf-488⁺ cells was quantified by flow cytometry. Statistical analysis was performed using multiple unpaired t-tests with false discovery rate correction. **(D)** Tf-488 recycling assays in A549 cells transfected with siRNA CTRL, siRNA USP7, or siRNA TRIM27. Cells were incubated with 15 µg/mL Tf-488 for 30 min at 4 °C, shifted to 37 °C for 1 h, washed, and incubated with unlabeled transferrin before fixation at the indicated time points to quantify the percentage of Tf-488⁺ cells by flow cytometry. Statistical analysis was performed using simple linear regression analysis. **(E)** DQ-OVA assays in A549 cells transfected with siRNA CTRL, siRNA USP7, or siRNA TRIM27. Cells were incubated with the indicated concentrations of DQ-OVA for 15 min at 37 °C, washed, and fixed to quantify the percentage of cells positive for DQ-OVA fluorescence (degraded OVA) by flow cytometry. Statistical analysis was performed using multiple unpaired t-tests with false discovery rate correction. **(F)** Same experiments as in Figure 5E, showing the median fluorescence intensity of DQ-OVA⁺ A549 cells at the indicated DQ-OVA concentrations. Statistical analysis was performed using multiple unpaired t-tests with false discovery rate correction. **(G)** Immunoblot showing EGFR degradation kinetics in A549 cells transfected with siRNA CTRL, siRNA USP7, or siRNA TRIM27 and treated with 100 ng/mL EGF, harvested at the indicated time points. Representative immunoblot from three independent experiments. **(H)** Quantification of EGFR signal from the experiments shown in Figure 5G. Statistical analysis was performed using simple linear regression. Data information: (A-F and H) Mean ± SEM of three biological replicates. p-values are denoted as follows: ns, not significant, *p < 0.05, **p < 0.01, ***p < 0.001, ****p < 0.0001.

### USP7 and TRIM27 depletion alter the endosomal fate of surface receptors and soluble cargos

In order to determine whether USP7 or TRIM27 silencing broadly enhances endocytic uptake or instead affects post-internalization steps of endosomal trafficking, we monitored the endosomal fate of transferrin, a classical marker used to study clathrin-mediated endocytosis and recycling. Transferrin maintains iron homeostasis by binding extracellular iron, undergoing transferrin-receptor-mediated endocytosis, releasing iron in endosomes, and recycling back to the plasma membrane (*49*). Of note, the transferrin receptor is only marginally degraded upon endocytosis, because its cytoplasmic domains contain recycling signals that direct it back to the plasma membrane for repeated cycles of transferrin uptake (*50*). Transferrin uptake assays were used to determine whether USP7 or TRIM27 depletion affects substrate uptake or receptor internalization. Control, USP7-depleted, and TRIM27-depleted A549 cells were incubated with fluorophore-labeled transferrin at 4 °C for 30 min to allow binding to the transferrin receptor, washed, and then shifted to 37 °C for 10 min to permit transferrin internalization. USP7 or TRIM27 knockdown did not increase transferrin uptake and even tended to reduce it, although these effects did not reach statistical significance (Fig. 5C). This trend is consistent with previous studies showing that impaired endosomal recycling can negatively impact endocytic uptake rather than enhance it (*51*). A similar workflow was used to perform transferrin recycling assays. After the binding step, cells were pulsed with fluorophore-labeled transferrin by shifting them to 37 °C, washed extensively, and then chased in medium containing excess unlabeled transferrin. We quantified the percentage of transferrin-positive cells at multiple time points over two hours to monitor transferrin recycling. A 10-min short pulse favors the fast-recycling route of transferrin, which can return to the plasma membrane while largely bypassing sorting endosomes. Under these conditions, 90% of cells recycled the transferrin input in less than 45 min, and USP7 or TRIM27 depletion did not alter these kinetics (fig. S4B). After a longer one hour transferrin pulse, internalized transferrin reaches sorting and recycling endosomes, and recycling proceeds through multiple Rab-dependent pathways, including fast recycling from early endosomes and slower recycling via the perinuclear recycling endosome (*49*, *52*). Consistent with this, recycling was slower under these conditions, and one hour was required for control siRNA cells to recycle 90% of the labelled transferrin (Fig. 5D). In contrast, USP7 or TRIM27 knockdown significantly delayed transferrin recycling after this long pulse, such that 90% of the labelled transferrin was recycled only after 90 min in depleted cells (Fig.5D). This delay matches that observed in Rab35-depleted cells, in which transferrin recycling is also significantly impaired (fig. S4C). Interestingly, Rab11 depletion did not reduce transferrin recycling under these conditions and even modestly increased it (fig. S4C). Together, these results suggest that USP7 and TRIM27 depletion does not broadly affect cargo uptake or the fast recycling of the transferrin-transferrin receptor complex but instead impairs the recycling of cargos that reach downstream sorting/recycling compartments.

Endosomal recycling defects are often associated with increased trafficking of soluble endosomal content to lysosomal degradative compartments (*51*, *53*, *54*). To test whether USP7 or TRIM27 depletion promotes such rerouting of the endosomal flow, we incubated USP7- and TRIM27-depleted A549 cells with increasing concentrations of DQ-OVA, a soluble ovalbumin derivative conjugated to a quenched fluorophore that becomes fluorescent upon proteolytic degradation. In this assay, DQ-OVA fluorescence reports on the functionality and magnitude of the endosomal-lysosomal degradation flux. Interestingly, USP7 or TRIM27 depletion enhanced DQ-OVA degradation, as a higher percentage of cells were DQ-OVA-positive, and the median fluorescence intensity of positive cells was significantly increased compared with control cells (Fig. 5, E and F). These experiments support a role for USP7 and TRIM27 in regulating endosomal trafficking by promoting endosomal recycling and preventing the routing of cargos towards lysosomal degradative compartments. A similar rerouting was observed in Rab35- and Rab11-depleted cells (Fig.S4D). Rab35 depletion promoted DQ-OVA degradation, whereas Rab11 knockdown tended to reduce it, consistently with their respective impact on transferrin recycling. Together, these findings reinforce a model in which endosomal recycling and degradation fluxes compensate for each other, such that impaired recycling favors soluble cargo degradation.

Our observations indicate that the USP7-TRIM27 complex is required to maintain endosome compartmentalization and endosomal trafficking, and that its depletion leads to enlarged EEA1⁺ early endosomes and remodeled LAMP1⁺ late endosomes (Fig. 3 and fig. S3). These functions are reminiscent of several well-characterized endosomal sorting and maturation factors (*55*). Notably, deficiencies in Rab11 and its effectors, or in ESCRT components involved in early-to-late endosome maturation, have been reported to create “traffic jams” in sorting endosomes, with transmembrane cargo accumulation in sorting and late endosomes, enlarged endosomal structures, and a collapse or remodeling of endosomal compartments (*47*, *51*, *56*, *57*). Consistent with this, the USP7-TRIM27 complex appears to act at the interface between early endosomal sorting and recycling, where it preserves endosomal compartment identity and balances recycling versus lysosomal degradation. In this model, USP7 or TRIM27 depletion would initially increase routing of endosomal flux toward LAMP1⁺ compartments but ultimately stall endosome trafficking of transmembrane cargo, causing endosomal “traffic jams,” impaired cargo delivery to degradative lysosomes, and delayed receptor downregulation. To test this, we treated USP7-, TRIM27-, or control siRNA-transfected cells with epidermal growth factor (EGF) and monitored EGFR degradation kinetics by western blot over time (Fig. 5, G and H). As expected, USP7 or TRIM27 silencing significantly delayed EGFR degradation, consistent with a defect in trafficking of transmembrane cargo to lysosomes. Altogether, these data support a fundamental role for the USP7-TRIM27 complex in maintaining the fidelity of endosomal sorting and recycling, analogous to factors in the Rab-dependent recycling pathways that also restrict ZIKV infection. Because depletion of the retromer components VPS35 and SNX17 failed to enhance ZIKV infection in our assays, we favor a model in which the USP7-TRIM27 complex associates with alternative endosomal sorting machineries, rather than acting solely through the canonical retromer pathway.

### Pathogenic USP7 mutations associated with Hao-Fountain syndrome promote ZIKV infection

Pathogenic variants in USP7 cause Hao-Fountain syndrome, a rare haploinsufficient neurodevelopmental disorder characterized by developmental delay, autism spectrum disorder, brain MRI anomalies, seizures, and other systemic features (*17*). USP7 haploinsufficiency arises from microdeletions or heterozygous loss-of-function variants, and emerging evidence also implicates gain- or altered-function missense variants in Hao-Fountain syndrome. Because USP7 regulates the stability and activity of numerous substrates, pleiotropic disruption of these pathways likely contributes to the broad clinical spectrum observed in affected individuals. To date, more than 250 families have been reported with pathogenic USP7 variants or USP7-containing microdeletions, but case identification still largely depends on access to genome or exome sequencing, and therefore the true prevalence remains unknown (*17*). In parallel, large-scale population databases such as gnomAD, which aggregates sequencing data from >800,000 individuals, list numerous USP7 missense variants of unknown significance. To prioritize potentially deleterious variants, we ranked USP7 missense mutations by their Combined Annotation-Dependent Depletion (CADD) score, a quantitative estimate of the predicted functional impact of a variant, irrespective of whether it has already been classified as pathogenic. A high CADD score is expected to indicate that a variant is functionally deleterious. We then applied the gene-specific mutation significance cutoff at the 95% confidence level (MSC95%), which defines a CADD threshold above which USP7 variants are considered among the most deleterious for this gene. For USP7, the MSC95% corresponds to a CADD score of 21.1. Using this threshold, we identified >600 USP7 missense variants in gnomAD with CADD scores above the MSC95%, which we consider candidate pathogenic alleles for Hao-Fountain syndrome. Their allele frequencies ranged from approximately 10^−5^ to 10^−7^, consistent with Hao-Fountain syndrome being categorized as an ultra-rare genetic disorder (Fig. 6A). By contrast, USP7 missense variants with higher allele frequencies generally had much lower CADD scores and are therefore more likely to represent benign variation. Among the highly disruptive USP7 missense variants identified in gnomAD, 7 have already been reported in patients with Hao-Fountain syndrome, supporting that USP7 variants with high CADD scores are *bona fide* candidate pathogenic mutations.

**Fig. 6.**
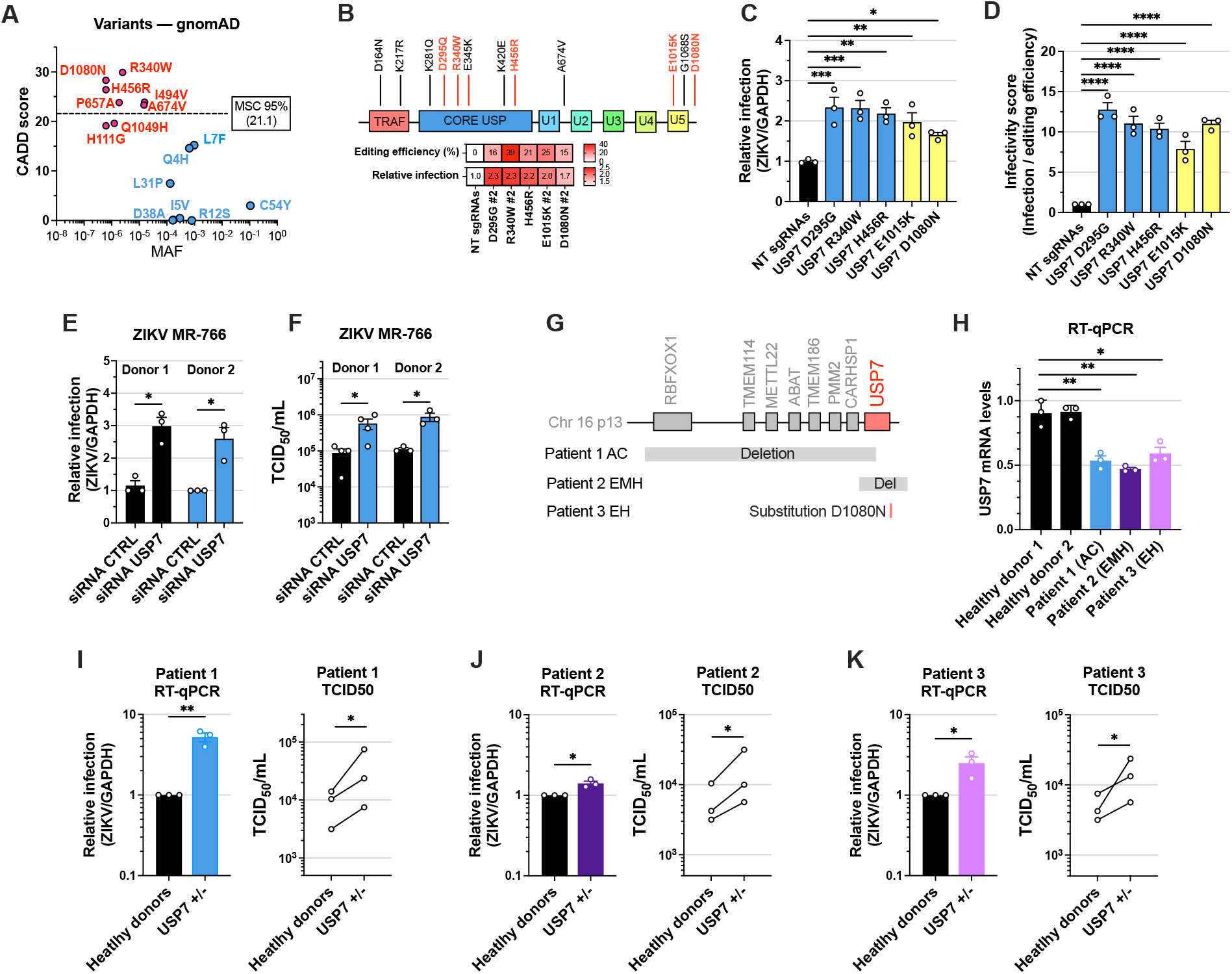
USP7 pathogenic mutations associated with Hao-Fountain syndrome promote ZIKV infection in A549 cells and human primary fibroblasts. **(A)** Analysis of USP7 variants from the gnomAD database. A selection of USP7 variants is shown according to their Combined Annotation–Dependent Depletion (CADD) score and minor allele frequency (MAF). Pathogenic USP7 missense variants reported in patients with Hao-Fountain syndrome are shown in red, and high-frequency USP7 missense alleles with likely benign effects are shown in blue. **(B)** Top: Selection of USP7 variants associated with Hao-Fountain syndrome that were introduced into A549 cells using CRISPR base editing. Variants shown in red were successfully edited. “Editing efficiency”: heatmap showing the percentage of editing for each variant 4 weeks post-transduction in A549 cells expressing the base editor and the corresponding sgRNA. “Relative infection”: heatmap showing the fold change in ZIKV infection for each edited A549 cell population. **(C)** Zika virus (ZIKV) strain MR-766 infection at 24 h post-infection in A549 cells expressing Cas9 base editors and sgRNAs introducing the indicated mutations. Infection was measured by RT-qPCR and normalized to GAPDH mRNA levels. Statistical analysis was performed using one-way ANOVA with Dunnett’s test. **(D)** “Infectivity score” associated with each USP7 pathogenic mutant introduced into A549 cells by CRISPR base editing. For each condition, the ZIKV infection rate was normalized to the corresponding editing efficiency to obtain a score reflecting the impact of each mutation on USP7 antiviral activity. Statistical analysis was performed using one-way ANOVA with Dunnett’s test. **(E)** ZIKV strain MR-766 infection at 24 h post-infection in human primary fibroblasts from two donors transfected with control siRNA pools (siRNA CTRL) or siRNA pools targeting USP7 (siRNA USP7). Infection was assessed by TCID50 titration of cell culture supernatants from cells infected at an MOI of 0.1. Statistical analysis was performed using multiple unpaired t-tests with false discovery rate correction. **(F)** USP7 silencing efficiency in human primary fibroblasts from two donors measured by RT-qPCR. Two-way ANOVA with Tukey’s test. **(G)** Schematic depicting the genotypes of the three Hao-Fountain syndrome patient donors. Patient 1 (AC) carries a large 16p13.2 deletion affecting USP7 and several upstream genes. Patient 2 (EMH) has a smaller deletion involving USP7. Patient 3 (EH) is heterozygous for the pathogenic USP7 missense variant D1080N. **(H)** USP7 transcript levels in human primary fibroblasts from USP7+/− donors, measured by RT-qPCR. Unpaired t-test. **(I)** ZIKV strain MR-766 infection at 24 h post-infection in human primary fibroblasts from USP7+/− patient 1. Infection was assessed by RT-qPCR and by TCID50 titration of cell culture supernatants from cells infected at an MOI of 0.1. An unpaired t-test was used for infection rates measured by RT-qPCR, and a paired t-test on log1010-transformed values was used for TCID50 titers. **(J)** ZIKV strain MR-766 infection at 24 h post-infection in human primary fibroblasts from USP7+/− patient 2. Infection was assessed by RT-qPCR and by TCID50 titration of cell culture supernatants from cells infected at an MOI of 0.1. An unpaired t-test was used for infection rates measured by RT-qPCR, and a paired t-test on log1010-transformed values was used for TCID50 titers. **(K)** ZIKV strain MR-766 infection at 24 h post-infection in human primary fibroblasts from USP7+/− patient 3. Infection was assessed by RT-qPCR and by TCID50 titration of cell culture supernatants from cells infected at an MOI of 0.1. An unpaired t-test was used for infection rates measured by RT-qPCR, and a paired t-test on log1010-transformed values was used for TCID50 titers. Data information: **(C-F, H-K)** Mean ± SEM of three biological replicates. p-values are denoted as follows: ns, not significant, *p < 0.05, **p < 0.01, ***p < 0.001, ****p < 0.0001.

Recent studies have reported pathogenic USP7 variants in individuals with Hao-Fountain syndrome. Building on these reports, we sought to introduce reported pathogenic USP7 missense mutations using CRISPR/Cas9 base-editing to define their impact on the antiviral function of USP7. Base-editing relies on a DNA deaminase fused to a catalytically inactive Cas9 to introduce C-to-T or A-to-G edits within a defined activity window, depending on the deaminase used (*58*). We designed 15 sgRNAs to introduce 12 patient-derived pathogenic missense variants across the USP7 coding sequence, together with one sgRNA designed to introduce a H456R substitution that targets a critical catalytic residue and is expected to disrupt USP7 enzymatic activity (Fig. 6B). Two of these variants target the N-terminal TRAF domain, which mediates protein-protein interactions, six target the catalytic core domain, one is located between the first and second UBL domains, and three others target to the C-terminal fifth UBL domain. A549 cells were transduced with lentiviral vectors expressing the Cas9-NG variant, which has a relaxed PAM requirement, fused either to the CBE4max cytidine deaminase or the ABE8e adenine deaminase, together with sgRNAs targeting the pathogenic sites or non-targeting sgRNAs as negative controls (*59*). Four weeks post-transduction, edited cell populations were challenged with ZIKV to assess the impact of USP7 pathogenic variants on its antiviral activity. Five mutations: D295G, E345K, and H456R in the catalytic domain, and E1015K and D1080N in the UBL5 domain, significantly enhanced ZIKV infection by approximately 50 to 200% (Fig 6, B and C and fig. S5A). To quantify editing efficiency, genomic DNA was extracted from transduced cells, the targeted USP7 loci were PCR-amplified, and Sanger chromatograms were analyzed with EditR to measure peak overlaps (*60*) (fig. S5B). For the five edits that increased ZIKV infection, editing efficiencies ranged from 15% to 40%, indicating successful, although low-to-moderate, editing at the population level. No unintended bystander mutations were detected for four of these edits; the H456R variant, however, was co-introduced with an S457G change at ∼20% editing efficiency (Fig. S5B). Notably, conditions that did not alter ZIKV infection corresponded either to unsuccessful editing or to edits dominated by bystander substitutions with unknown effects on USP7 function. Although overall editing efficiencies were modest, these experiments demonstrate that pathogenic USP7 missense mutations can promote ZIKV infection. To estimate the relative impact of each variant on USP7 antiviral activity, we combined the observed fold change in ZIKV infection with the corresponding editing efficiency to derive an “infectivity score” for each successfully edited population (Fig. 6D). All five pathogenic variants that enhanced ZIKV infection yielded comparable infectivity scores between 7 and 10, indicating that, among the variants we were able to introduce, no major differences were detected in their effect on USP7 antiviral function. This suggests that pathogenic missense mutations associated with Hao-Fountain syndrome may impair the antiviral activity of USP7 to a similar extent, although further data will be required to determine whether this is a general property of USP7 pathogenic alleles.

### Hao-Fountain patient-derived fibroblasts are more susceptible to ZIKV infection

After delivery into the skin by an infected mosquito, ZIKV infects epidermal keratinocytes and dermal fibroblasts, which support viral replication and amplify viral load locally before the virus disseminates to distal organs (*61*, *62*). Hence, we sought to determine whether USP7 limits ZIKV infection in primary fibroblasts, which represent biologically relevant target cells *in vivo*. Human primary fibroblasts from two unrelated healthy donors were transfected with siRNA pools targeting USP7 or with non-targeting control siRNA pools prior to ZIKV infection (Fig. 6, E and F and fig S5C). Consistent with our results in cell lines, USP7 depletion in primary fibroblasts from both donors increased ZIKV RNA replication by 2 to 3-fold and viral production by 5- to 10-fold, which support an antiviral role for USP7 in orthoflavivirus target cells (Fig. 6, E and F). siRNA transfection efficiently silenced USP7 expression, reducing USP7 transcript levels by approximately 85% (fig S5C).

We next investigated whether primary fibroblasts derived from individuals with Hao-Fountain syndrome exhibit increased susceptibility to ZIKV infection. Three patient-derived fibroblast lines were obtained from the CombinedBrain biorepository (Figure 6G). Primary fibroblasts from patient 1 harbor a large chromosome 16 deletion resulting in the loss of *USP7* together with *CARHSP1*, *PMM2*, *TMEM186*, *ABAT*, *METTL22*, *TMEM114*, and RBFOX1, which are located in the same chromosomal region. Fibroblasts from patient 2 carry a smaller deletion affecting USP7. Fibroblasts from patient 3 express the pathogenic USP7 D1080N substitution, for which we previously showed by CRISPR/Cas9 base-editing that it impairs USP7 antiviral function (Fig. 6, B and C). All three patient-derived fibroblast lines expressed significantly lower USP7 transcript levels than fibroblasts from healthy donors (Fig. 6H). Although USP7 expression in the D1080N line was slightly higher than in the deletion cells, it remained reduced compared with controls, which may reflect donor-specific variation or additional, uncharacterized genetic changes. Patient fibroblasts were challenged with ZIKV, and infection levels were quantified by RT-qPCR and TCID50 assays in parallel with fibroblasts from two healthy donors (Fig. 6, I to K). Fibroblasts derived from patients 1 and 3 displayed significantly higher ZIKV infection, both at the level of viral RNA and infectious viral production, with patient 1 showing the largest increase (approximately 5-fold). For patient 2, viral RNA levels were only slightly elevated compared with controls, but infectious viral production was increased by about 2-fold (Fig. 6J). Together, these infection assays indicate that Hao-Fountain syndrome patient-derived fibroblasts are more susceptible to ZIKV infection, supporting our observations from cell lines and primary fibroblasts. Altogether, our results show that USP7 exerts antiviral activity against orthoflaviviruses primary human cells, and phenotypes from patient-derived fibroblasts raise the possibility that USP7 mutations could underlie inborn errors of immunity affecting antiviral defense.

### Anecdotal reports of recurrent respiratory infections in Hao-Fountain syndrome align with USP7 antiviral activity against SARS-CoV-2

Although Hao-Fountain syndrome has multisystem manifestations, early clinical descriptions did not identify an immunophenotype (*16*, *17*, *63*). However, recurrent respiratory infections from early childhood have been reported in three isolated cases (*18–20*). Two involved individuals have pathogenic *USP7* variants, whereas the third has a co-occurring *CFTR* variant that likely accounted for the respiratory phenotype. Because USP7 restricts orthoflavivirus entry through endocytosis, we asked whether it similarly limits respiratory viruses that use this route. In TMPRSS2-negative A549-ACE2 cells, where SARS-CoV-2 WA1 enters predominantly through endocytosis depletion of either USP7 or TRIM27 significantly increased infection by 3 to 10-fold (Fig. S5C). These findings extend the antiviral activity of the USP7-TRIM27 axis beyond orthoflaviviruses and support the possibility that USP7 deficiency contributes to an inborn error of immunity.

## DISCUSSION

Pathogenic variants in multifunctional genes can cause inborn errors of immunity by disrupting intrinsic antiviral defenses, innate immune signaling, or virus-induced cell death. Using an arrayed siRNA screen with ZIKV as a model virus, we tested NDD-associated genes with reported functions overlapping the orthoflavivirus life cycle. A subset significantly affected ZIKV infection, IFN responses, and inflammatory signaling. The strongest antiviral hit, NECAP1, regulates clathrin-mediated endocytosis; its depletion likely enhances viral entry by increasing uptake. Pathogenic NECAP1 variants are associated with epileptic encephalopathy, severe and frequent seizures, developmental and motor delay, and other brain abnormalities (*25*). Notably, a 7-year-old boy from Saudi Arabia with a homozygous premature-stop NECAP1 variant died in the setting of uncontrolled seizures and a lower respiratory tract infection caused by an unidentified pathogen (*24*). Given the central role of clathrin-mediated entry in infection by many respiratory viruses, impaired NECAP1 function could promote viral uptake, high viral loads, and severe infectious disease. This model parallels observations for TMEFF1, which regulates HSV-1 internalization and whose pathogenic variants predispose to severe HSV-1 encephalitis (*64*, *65*).

Our screen also identified YIPF5, an ER-resident factor, as a ZIKV antiviral gene that limits infection-induced IL-6 transcription. YIPF5 is linked to ER stress and a syndromic disorder characterized by microcephaly, epilepsy, and early-onset diabetes (*28*, *29*). Because YIPF5 regulates ER-to-Golgi transport, its depletion impairs DNA-virus sensing and downstream immune responses by preventing appropriate STING trafficking to the Golgi and limiting TBK1 signaling. Its immune functions are particularly relevant to ZIKV infection, as YIPF5-associated pathways appear to restrict ZIKV infection and inflammation while promoting neurogenesis and brain development processes central to congenital ZIKV pathogenesis.

We focused on USP7 and defined its previously unrecognized cell-intrinsic antiviral activity against selected neurotropic orthoflaviviruses. ZIKV and WNV were strongly restricted by the USP7-TRIM27 complex, which regulates endosomal trafficking and appears to limit virion internalization into target cells. We extended this observation to other endosomal-recycling regulators: depletion of Rab11, Rab4, or Rab35 markedly enhanced ZIKV infection, indicating that ZIKV is highly sensitive to endosomal-recycling function. This sensitivity was selective, as depletion of other recycling-pathway components, including the retromer subunit VPS35 and SNX17, did not measurably affect ZIKV infection. Moreover, DENV-2 16681 and YFV 17D were unaffected by loss of USP7, TRIM27, or Rab11. These findings reveal unappreciated differences in how orthoflaviviruses engage endocytic routes and traffic through endosomal compartments and suggest that some viruses may evade endocytic pathways that limit entry. Future studies should define the viral and host determinants governing sensitivity to these pathways and relate these differences to the marked variation in orthoflavivirus tissue tropism and dissemination.

Our data establish a role for the USP7-TRIM27 axis in endosomal recycling and endosomal compartmentalization but do not yet resolve how these pathways restrict infection. Recycling may directly prevent entry by returning internalized virions to the extracellular milieu, or its disruption may instead favor transport toward late, acidic, fusion-competent endosomes. Although both models remain possible, ZIKV internalization assays favor active restriction of virion access to the intracellular milieu by USP7 and TRIM27. Their depletion did not increase bulk endocytic uptake in transferrin-uptake assays. USP7 depletion also had a stronger effect than TRIM27 depletion on endosomal organization, including the accumulation of large, apparently empty vesicles, whereas TRIM27 depletion produced a 2-to 5-fold stronger antiviral phenotype. These findings suggest that USP7 and TRIM27 functions partially overlap, and that disruption of the endosomal network alone does not determine the magnitude of antiviral restriction.

We also did not assess whether USP7 or TRIM27 depletion alters the expression or localization of WNV and ZIKV receptors. However, no receptor has been shown to be uniquely required for WNV and ZIKV entry without also affecting DENV-2 or YFV infection. This specificity, together with evidence that impaired endosomal recycling generally does not increase receptor delivery to the plasma membrane, and can reduce surface receptor abundance and endocytosis, argues against altered receptor abundance as the principal explanation for our phenotype. Nevertheless, unknown WNV- and ZIKV-specific receptors, differences in envelope-receptor affinity, and distinct sensitivity to pH-dependent envelope conformational changes cannot be excluded. Higher-resolution particle-tracking approaches will be necessary to determine whether ZIKV and WNV are actively returned to the cell surface through recycling pathways, as confocal imaging of incoming virions using anti-E staining lacked sufficient resolution to visualize these particles.

More broadly, our findings suggest that endosomal recycling can function as an antiviral barrier by preventing virions from reaching fusion-competent endosomal membranes. This poorly characterized form of cell-intrinsic defense may be especially relevant in genetic and chronic disorders affecting trafficking pathways. Flux-based cellular systems are particularly vulnerable to disruption, as impairment of a single component can cause widespread functional consequences. In addition to USP7-associated Hao-Fountain syndrome, impaired endosomal recycling occurs in disorders involving related complexes, including MAGEL2-associated Prader-Willi and Schaaf-Yang syndromes and diseases caused by WASH-complex mutations. Endosomal trafficking defects and neurotoxicity have also been implicated in Alzheimer’s and Parkinson’s diseases. Framing endosomal recycling as an immune pathway may therefore help uncover the molecular basis of established comorbidities that increase susceptibility to, or severity of, infectious diseases.

## MATERIAL AND METHODS

### Plasmids

pLX_311-Cas9 was a gift from Prof J. Doench (Addgene Plasmid #96924) (*66*). Lentiviral vectors coding for sgRNAs were obtained by cloning annealed oligonucleotides in BsmBI-digested pLentiGuide-Hygro, a gift from Prof C. Goujon (Addgene Plasmid #139462). pRDA_78_Cas9_NG was obtained by cloning the BE4max-spCas9-NG coding sequence from pCAG-CBE4max-SpCas9-NG-P2A-EGFP (Addgene Plasmid #140001) (*59*) in pRDA_78 (Addgene Plasmid #158582)(*67*) by Gibson assembly (New England Biolabs HiFi DNA Assembly kit), using the AgeI and Bsu36I restriction sites for vector linearization. pRDA_429 was a gift from Prof J. Doench (Addgene Plasmid #179098).

pRRL_SFFV_3xFLAG_USP7_IRES_Puro, pRRL_SFFV_3xFLAG_eGFP_IRES_Puro, pRRL_SFFV_HA_TRIM27_IRES_Puro, and pRRL_SFFV_3xFLAG_TRIM27_IRES_Puro were generated by cloning synthetic DNA fragments encoding 3xFLAG-USP7, 3xFLAG-eGFP, or HA-TRIM27 (Twist Bioscience) into BamHI/XhoI-digested pRRL_SFFV_E2Crimson_IRES_Puro, a gift from C. Goujon (Addgene #139445). pCDNA4_3xFLAG-USP7 was obtained by subcloning 3xFLAG-USP7 from pRRL_SFFV_3xFLAG_USP7_IRES_Puro into BamHI/XhoI-digested pCDNA4, and pCDNA4_3xFLAG-eGFP was generated by subcloning 3xFLAG-eGFP from pRRL_SFFV_3xFLAG_eGFP_IRES_Puro into BamHI/XhoI-digested pCDNA4. Primers and oligonucleotide sequences used for molecular cloning are listed in Table S2.

### Cell lines and primary cells

Human cell lines HEK293T (ATCC CRL-3216) and Vero CCL-81 were obtained from the ATCC (American Type Culture Collection). The A549-ACE2 cell line was a kind gift from Brad Rosenberg (Icahn School of Medicine at Mount Sinai, NY, USA) and is referred to as A549 throughout this manuscript. Huh7.5 cell line was a kind gift from Matt Evans (UC Irvine, CA, USA). Healthy primary fibroblasts and Hao-Fountain patient fibroblasts were obtained from CombinedBrain. Vero E6 (ATCC, CRL-1586) modified to express TMPRSS2 were a kind gift from Adolfo Garcia-Sastre (Icahn School of Medicine at Mount Sinai, New York, NY). A549-ACE2-Cas9 were obtained by transduction of A549-ACE2 with HIV-1-based lentiviral vectors expressing the spCas9 (Addgene plasmid # 96924). A549-ACE2-Cas9 cells were generated by transducing A549-ACE2 cells with an HIV-1–based lentiviral vector expressing spCas9 (Addgene #96924). A549-ACE2 cells stably expressing the CBE4max-SpCas9-NG or ABE8e-SpCas9-NG base editors were generated by transduction with HIV-1–based lentiviral vectors derived from pRDA_78_Cas9_NG or from pRDA_429 (Addgene #179098), respectively. All cell lines were maintained at 37 °C in a humidified incubator with 5% CO₂ in Dulbecco’s modified Eagle medium (DMEM, Corning) supplemented with 10% (vol/vol) fetal bovine serum (FBS, Thermo Fisher Scientific) and 100 U/mL penicillin plus 100 µg/mL streptomycin (Corning). For antibiotic selection, cells were treated with 10 μg/ml Blasticidin (Fisher), 1μg/ml Puromycin (Fisher), 100μg/ml Puromycin (Invivogen). Human primary fibroblasts were cultured in Alpha MEM Eagle (PAN-Biotech) supplemented with 10% FBS and penicillin–streptomycin. All cultures were tested monthly and confirmed negative for mycoplasma contamination (Lonza).

### siRNA screen and siRNA transfection

Knockdown of the indicated targets was achieved by reverse transfection of siRNA pools or individual siRNAs (siGENOME, Horizon Discovery) using Lipofectamine RNAiMAX (Thermo Fisher Scientific) according to the manufacturer’s instructions. A final siRNA concentration of 44 nM was used in 12-well plates. For the screen shown in Figure 1, custom siGENOME siRNA pools, including two non-targeting control pools, were supplied pre-arrayed in 96-well plates (Horizon Discovery). siRNAs were resuspended to 10 µM in nuclease-free water and reverse-transfected into A549 cells as indicated. Three days after transfection, cells were infected with ZIKV strain MR-766 for 24 h, after which culture supernatants were harvested for TCID50 titrations on Vero CCL-81 cells and infected cells were lysed for RNA extraction and RT-qPCR analysis. The siRNA targets used are listed in Table S6.

### CRISPR/Cas9 knock-out

For CRISPR/Cas9 knock-out in A549 lines, lentiguide-hygro lentiviral vectors coding sgRNAs targeting the indicated genes or non-targeting sgRNAs were produced, and A549-ACE2-spCas9 cells were transduced for 6 h before replacing the supernatants with fresh, complete medium. The transduced cells were selected with hygromycin 2 days later and amplified for 12–15 days before infection assays and western-blotting.

### CRISPR/Cas9 base editing

For base-editing experiments in A549-derived cell lines, lentiviral vectors encoding Cas9-NG fused to the appropriate base editor together with sgRNAs targeting the indicated loci or non-targeting control sgRNAs were produced and used to transduce A549-ACE2 cells for 6 h, after which supernatants were replaced with fresh complete medium. Transduced cells were selected with puromycin for 2 days and then maintained for 4 weeks prior to infection assays and genomic DNA extraction. Genomic DNA was amplified by PCR, and amplicons were subjected to Sanger sequencing to determine editing efficiency. Editing efficiencies were quantified from sequencing chromatograms using the EditR online tool, which measures base-editing frequencies from peak intensities and overlaps. Oligonucleotides encoding the sgRNAs are listed in Table S3, and primers used for PCR on genomic DNA and for Sanger sequencing are listed in Table S4.

### Orthoflavivirus production and infection

ZIKV strain MR-766 and WNV strain NY99 were kindly provided by J. Lim (Icahn School of Medicine at Mount Sinai, New York, USA) and obtained from BEI Resources (NR-50065 and NR-158, respectively). DENV-2 strain 16681 was kindly provided by M. Evans (Icahn School of Medicine at Mount Sinai, New York, USA), and YFV strain 17D was provided by C. Rice (Rockefeller University, New York, USA). DENV-2 16681 stocks were generated by transfecting 293T cells with a plasmid encoding this viral cDNA under the control of a polymerase II promoter (manuscript in preparation) using a protocol we previously described (*70*). Approximately 4×10^5^ 293T cells per well were seeded into poly-lysine coated 6-well plates. The following day, immediately prior to transfection the media was changed to 1mL of 3% DMEM, and cells were transfected with 2 µg of DENV plasmid per well and TransIT-LT1 transfection reagent (Mirus Bio, Madison, WI) per the manufacturer’s recommendations. Six hours post transfection, the media was replaced with 1.5mL of 3% DMEM. Supernatants were collected on day 3 and 4 post transfection, pooled, centrifuged to remove cellular debris, and stored at -80°C. YFV strain 17D was generated from an infectious clone plasmid encoding the full-length genome by linearization and *in vitro* transcription using a mMessage mMachine T7 transcription kit (Ambion). Huh-7.5 cells were electroporated with viral RNA and incubated at 37°C. Supernatants containing progeny virions were collected from electroporated cells on day 2 post-electroporation, clarified by centrifugation, and stored at -80°C. Working stocks for all viruses were prepared by amplification on Vero CCL-81 cells (ATCC CCL-81) using a MOI of 0.001 in DMEM containing 2% FCS. After 3-5 days, upon appearance of cytopathic effects, culture supernatants were harvested, clarified by centrifugation to remove cell debris, aliquoted, and stored at -80 °C. Viral titers were determined by TCID50 assay on Vero CCL-81 cells. Cells were infected at the indicated multiplicity of infection (MOI, calculated from titers obtained in Vero CCL-81 cells) by incubating cells with viral inputs in serum-free medium for 1 h, washing once with PBS, and then adding serum-containing medium for the indicated times. When indicated, cells were pretreated overnight with a combination of TBK1/IKKε inhibitor (InvivoGen) and pan-JAK inhibitor (Sigma-Aldrich) at 500 nM each, or for 1 h with Pitstop 2 (MedChemExpress) at 1 µg/mL, and the compounds were maintained at the same concentrations throughout the course of infection.

### SARS-CoV-2 production and infection

SARS-CoV-2 isolate USA-WA1/2020 (NR-52281) was obtained from BEI Resources, NIAID, NIH. Virus stocks were amplified on Vero E6 TMPRSS2 cells in DMEM 2% FBS using an MOI of 0.001. Supernatant was collected 2-3 days post-infection when cytopathogenic effects started to be observed, cleared by centrifugation and aliquots were frozen down at -80 °C. Virus stock titers were determined by plaque assay on Vero E6 cells. All work with live virus was done in the CDC/USDA-approved biosafety level 3 (BSL-3) facility of the Icahn School of Medicine at Mount Sinai in accordance with their respective guidelines for BSL-3 work. Cells were infected at the indicated MOI by incubating cells with viral inputs in DMEM 2% FBS for the indicated times.

### TCID50/mL titration

Infectious virus in cell culture supernatants was quantified by TCID50/mL titration on Vero CCL-81 cells seeded at 20,000 cells per well in 96-well plates. Twenty-four hours after plating, cells were inoculated with 100 µL of 10-fold serial dilutions of supernatants in DMEM containing 2% FCS, using 8 replicate wells per dilution. After 5–7 days of incubation at 37 °C, wells were examined for cytopathic effect, and virus titers were calculated using the Reed– Muench method.

### RNA quantification by RT-qPCR

Target silencing efficiency and viral RNA levels were assessed by collecting cells 4 days after siRNA transfection or at the indicated times after infection, followed by isolation of total RNA using the RNeasy kit with on-column DNase treatment (Qiagen). cDNA was generated from 125–250 ng total RNA using the High-Capacity cDNA Reverse Transcription Kit (Thermo Fisher Scientific). Quantitative real-time PCR was performed in technical triplicates on a LightCycler 480 Instrument II (Roche) using SYBR Green PCR master mix (Life Technologies). For relative quantification, target mRNA or viral RNA levels were normalized to GAPDH mRNA expression and analyzed using the ΔΔCt method. Primer sequences are provided in Table 5.

### Single-round infectivity assay

Cells were infected with ZIKV strain MR-766 at the indicated MOI as described above. Four hours post-infection, the culture medium was either left unchanged or supplemented with 20 mM NH_4_Cl, and infection was allowed to proceed until 24 h post-infection. Viral replication/spread was quantified by RT-qPCR analysis of cell lysates at 24 h.

### Viral binding and internalization assay

For binding assays, cells were incubated with ZIKV strain MR-766 at MOI 10 at 4 °C for 1 h in 2% FCS medium. Cells were then washed extensively with PBS to remove unbound virus and lysed for RNA extraction and viral RNA quantification by RT-qPCR. For internalization assays, virus was first bound to cells at 4 °C as above, after which cultures were shifted to 37 °C for the indicated times to allow internalization. At each time point, cells were trypsinized to remove remaining surface-bound virus, washed with PBS, and lysed for RT-qPCR analysis. To verify that trypsinization efficiently removed non-internalized virus, cells were lysed immediately before the temperature shift (0 h time point). In the experiments shown in Fig. 3C, internalization data are plotted on the same scale as the binding assay, and viral RNA levels at 0 h were comparable to signal in mock-infected controls.

For internalization assays with pharmacological inhibitors, cells were pretreated for 1 h at 37 °C with 20 mM NH_4_Cl, 50 nM bafilomycin A1 (Sigma-Aldrich), 10 µg/mL cycloheximide (Sigma-Aldrich), or an equivalent volume of DMSO before performing the internalization experiments described above. The compounds were maintained at the same concentrations throughout the assay.

### Cell viability assays

Cell viability was assessed in siRNA-transfected cells by measuring cellular ATP levels in infected or non-infecting cells using the CellTiter-Glo® Luminescent Cell Viability Assay kit (Promega) according to manufacturer’s instructions.

### Confocal microscopy

siRNA-transfected, transduced, or unmodified cells were seeded at 70– 80% confluence in 24-well plates containing 12 mm #1.5H glass coverslips (Neurovitro Corporation # GG-12-1.5H) pre-coated with poly-L-lysine (Sigma-Aldrich). Twenty-four hours later, cells were fixed in 2-4% paraformaldehyde (PFA) in PBS for 15 min at room temperature, washed three times with PBS, and incubated with 50 mM NH_4_Cl in PBS for 10 min to quench free aldehydes. After additional PBS washes, cells were either permeabilized with 0.1% Triton X-100 in PBS for 15 min or with ice cold methanol for 10 min at -20C. Cells were then washed with PBS and blocked for 1h at RT in PBS 5% BSA 0.3% Triton X-100. Coverslips were then inverted onto 30 µL of primary antibody diluted to either 1:50, 1:100 or 1:300 in PBS 1% BSA 0.3% Triton X-100 and incubated overnight at 4°C in a humidified chamber. After incubation, coverslips were returned to the wells, washed several times with PBS, and incubated with fluorophore-conjugated secondary antibodies diluted 1:1,000 in PBS 1% BSA 0.3% Triton X-100 for 1h at room temperature in the dark. The following primary antibodies were used: anti-EEA1 (rabbit, Cell Signaling #2411), anti-USP7 (mouse, Thermo Fisher Scientific #MA5-31515, for colocalization with FLAG-TRIM27, or rabbit, Cell Signaling #4833), anti-LAMP1 (rabbit, Cell Signaling #9091), anti-FLAG (rabbit, Sigma-Aldrich #F7425), anti-TRIM27 (rabbit, Cell Signaling #15099), and ActinGreen 488 ReadyProbes phalloidin (Invitrogen #R37110). When phalloidin or other directly labeled probes were used together with unconjugated primary antibodies, directly labeled reagents were added after incubation with non-conjugated primaries for 30 min at room temperature in the dark. Secondary antibodies were goat anti-mouse IgG-Alexa Fluor 647 (Thermo Fisher #A-121235), goat anti-mouse Alexa Fluor 488 (Thermo Fisher #A-11001), goat anti-rabbit IgG–Alexa Fluor 568 (Thermo Fisher #A-11011) or goat anti-rabbit Alexa Fluor 647 (Thermo Fisher #A-21245). Finally, nuclei were counterstained with Hoescht 33342 (Thermo Fisher #H3570) at 1:2,000 in PBS, and coverslips were mounted in ProLong Glass Antifade Mountant (Thermo Fisher #P36982) on SuperFrost slides (Electron Microscopy Sciences #71867-01). After curing overnight in the dark, coverslips were sealed with clear nail polish, and slides were stored at −20 °C until imaging on a Zeiss LSM980 confocal microscope Airyscan 2 (Carl Zeiss). Images were acquired with a Plan-Apochromat 63x/1.4 by oil immersion and the operating software ZEN Blue. By default, a zoom of 1.5 was applied and for specified fields, an additional zoom was further performed, as indicated in the legend of the figure. Images were analyzed using Fiji (ImageJ).

### Electron microscopy

siRNA-transfected cells were seeded in permanox chamber slides (EMS, #70390) and fixed with 2% paraformaldehyde/ 2.5% glutaraldehyde in 0.1M sodium cacodylate solution (EMS, #15960-01) at 4°C before being embedded using standard transmission electron microscopy embedding protocols. Briefly, cells were rinsed in 0.1M sodium cacodylate buffer, post fixed with 2% osmium tetroxide/1.5% potassium ferricyanide in 0.1M sodium cacodylate, and *en bloc* stained with aqueous 2% uranyl acetate. Cells were dehydrated in an increasing concentration of ethanol (25% up to 100%), infiltrated with an ascending ethanol and an EPON resin series (Embed 812 Kit, EMS, 14120), and placed in pure resin overnight. Modified BEEM capsules (EMS, 69910-01) were placed on top of the cells, filled with pure resin, and heat polymerized at 60°C for 72 h. Post polymerization, capsules were snapped from the substrate to dislodge the cells. Samples were placed in a 60°C vacuum oven for 72 h to polymerize. Semithin sections (0.5 um) were obtained using a Leica UCT ultramicrotome (Leica, Buffalo Grove, IL) and counterstained with 1% toluidine blue. Ultra-thin sections (80nm) were collected onto copper 200 mesh grids (EMS, G200H-Cu) using a Coat-Quick adhesive pen (EMS, 70624). Sections imaged on an HT7500 transmission electron microscope (Hitachi High-Technologies, Tokyo, Japan) using an AMT NanoSprint12 12-megapixel CMOS TEM Camera (Advanced Microscopy Techniques, Danvers, MA). Images were only adjusted for contrast on the AMT software.

### Western Blotting

Cells were lysed in RIPA buffer containing SDS (50 mM Tris-HCl pH 7.4, 150 mM NaCl, 1% Triton X-100, 0.5% sodium deoxycholate, 0.1–1% SDS). Clarified lysates were mixed with Laemmli sample buffer (Bio-Rad) and 10% (vol/vol) β-mercaptoethanol, then boiled for 10 min before SDS–PAGE and transfer to nitrocellulose membranes (Bio-Rad). Membranes were blocked in 5% (wt/vol) non-fat dry milk in TBS-T for 1 h at room temperature. Blots were incubated overnight at 4 °C with primary antibodies diluted in 5% milk or 5% BSA in TBS-T as indicated: anti-USP7 (rabbit, Cell Signaling #4833, 1:1,000), anti-TRIM27 (rabbit, Thermo #12205-1-AP, 1:1,000), anti-GAPDH (rabbit, Cell Signaling #2118, 1:10,000), anti-HA (rabbit, Cell Signaling #3724, 1:10,000), anti-FLAG-HRP (mouse, Sigma #A8592, 1:10,000), and anti-EGFR (rabbit, Cell Signaling #4267, 1:1,000). Membranes were washed three times for 5 min each in TBS-T, then, if necessary, incubated with HRP-conjugated secondary antibody (anti-rabbit IgG– HRP, Cell Signaling #7074) diluted in TBS-T for 1 h at room temperature, followed by three additional 5-min washes in TBS-T. Signals were detected using ECL reagent (Thermo Fisher Scientific) and captured either on a chemiluminescence imager (Bio-Rad) or on Hyperfilm (Cytiva Life Sciences) using a film developer.

### Flow cytometry analysis of infected cells

A total of 30,000 cells per condition were infected in 96-well plate format at the indicated times and MOIs, then fixed in 4% paraformaldehyde for 15 min at room temperature. Cells were washed once with PBS and incubated twice for 15 min in intracellular staining buffer (PBS supplemented with 0.2% BSA and 0.05% saponin) prior to antibody staining. Cells were then incubated for 1 h at room temperature with the pan-flavivirus monoclonal antibody 4G2 (1:1,000; 1 mg/mL stock, a gift from Prof. F. Krammer, Icahn School of Medicine at Mount Sinai, New York, USA) or the anti-SARS-CoV-1/2 N 1C7C7 antibody (1:1,000; 1mg/mL stock) diluted in staining buffer. After three washes in staining buffer, cells were incubated for 1 h at room temperature with 1 µg/mL (1:1,000) goat anti-mouse IgG-PE secondary antibody (Thermo Fisher Scientific) in staining buffer, washed twice, and resuspended in PBS. E-positive cells were quantified on an Attune NxT flow cytometer (Thermo Fisher Scientific), and data were analyzed using FlowJo software (TreeStar, USA).

### Transferrin uptake and recycling assays

siRNA-transfected A549 cells were plated in 24-well format and used for assays the following day. Cells were starved for 1-2 h at 37°C in incubation buffer (DMEM supplemented with 0% FCS, 25 mM HEPES, and 0.2% BSA) prior to addition of Alexa Fluor 488-conjugated transferrin (Thermo Fisher). For uptake assays, cells were chilled to 4°C and incubated with the indicated concentrations of conjugated transferrin for 30 min at 4°C. Cells were then washed three times with PBS to remove unbound transferrin and transferred to 37°C in incubation buffer to allow synchronized internalization. After 15 min, internalization was stopped by washing cells with ice-cold PBS, and extracellular fluorescence from membrane-bound transferrin was removed by washing with an acid stripping solution (0.1 M glycine-HCl, 0.15 M NaCl, pH 2-3). Cells were then scraped, fixed in 4% paraformaldehyde for 15 min, and resuspended in PBS for flow cytometry analysis. For recycling assays, starved cells were incubated with 20 µg/mL Alexa Fluor 488-conjugated transferrin for 10 min or 1 h at 37°C, as indicated. Cells were then washed three times with PBS to remove unbound conjugated transferrin and incubated in complete DMEM supplemented with 100 µg/mL unlabeled transferrin (Fisher) to drive recycling. At the indicated time points, transferrin recycling was stopped by washing cells three times with ice-cold PBS. Cells were then detached by scraping in trypsin solution, fixed in 4% paraformaldehyde for 15 min, and resuspended in PBS for flow cytometry analysis.

### DQ-OVA proteolysis assays

A total of 120,000 siRNA-transfected A549 cells were plated in 24-well format and used for assays the following day. Cells were incubated with the indicated concentrations of DQ-OVA (Thermo Fisher) at 37°C for 30 min, washed with ice-cold PBS, scraped, fixed in 4% paraformaldehyde for 15 min, and resuspended in PBS for flow cytometry analysis. Analysis quantified the percentage of DQ-OVA-positive cells (reflecting proteolytic processing of DQ-OVA) and the median fluorescence intensity of DQ-OVA-positive cells.

### EGFR degradation assay

A total of 120,000 siRNA-transfected A549 cells were plated in 24-well format and starved in incubation buffer (DMEM supplemented with 0% FCS, 25 mM HEPES, and 0.2% BSA) for assays the following day. Cells were pre-treated with 10 µg/mL cycloheximide (Sigma-Aldrich) for 1 h before stimulation with 100 ng/mL EGF (R&D Systems) to trigger EGFR internalization. At the indicated time points, cells were washed with PBS and lysed in RIPA buffer for western blot analysis.

## ACKNOWLEDGMENTS

This work was supported by the National Institutes of Health through NIH R21AI187731 (J.R.J.) and R01AI124690 (C.M.R.). We thank the members of the Johnson, Lim, Rosenberg, Evans, García-Sastre, and Rice laboratories, as well as the ISMMS Department of Microbiology, for helpful feedback and support. We are grateful to Holly Ramage, Nolwenn Jouvenet, Maudry Laurent-Rolle, Scott B. Biering, Caroline Goujon, Olivier Moncorgé, Valérie Courgnaud, Maïka Deffieu, Sébastien Kury, Dusan Bogunovic, Maria Gabriela Noval, Nikolas Klink, and Malte Gersch for generously providing reagents and protocols and for helpful discussions. We thank Randy Albrecht, Lokendrasingh Chauhan, and Adam Abdeljawad for their support with the BSL-3 facility and procedures at the ISMMS. Confocal microscopy and electron microscopy sample preparation and imaging were performed at the Microscopy and Advanced Bioimaging Core at the ISMMS. We also thank the Foundation for Hao-Fountain Syndrome, Bo Bigelow, Amber N. Freed, Becky Raatz, and the broader Hao-Fountain syndrome community for their generous support. We thank the Manger la Vie Foundation; Olivier and Stéphanie Emmenecker; Cécile and Jean Brin; and the foundation community for their support. Finally, we thank the COMBINEDBrain team, biorepository teams, and anonymous donors for their generous support of this research.

## REFERENCES

1. S. J. Chapman, A. V. S. Hill, Human genetic susceptibility to infectious disease. Nat. Rev. Genet. 13, 175–188 (2012).

2. A. J. Kwok, A. Mentzer, J. C. Knight, Host genetics and infectious disease: new tools, insights and translational opportunities. Nat. Rev. Genet. 22, 137–153 (2021).

3. M. C. Poli, I. Aksentijevich, A. A. Bousfiha, C. Cunningham-Rundles, S. Hambleton, C. Klein, T. Morio, C. Picard, A. Puel, N. Rezaei, M. R. J. Seppänen, R. Somech, H. C. Su, K. E. Sullivan, T. R. Torgerson, I. Meyts, S. G. Tangye, Human inborn errors of immunity: 2024 update on the classification from the International Union of Immunological Societies Expert Committee. J. Hum. Immun. 1, e20250003 (2025).

4. Y. T. Akalu, D. Bogunovic, Inborn errors of immunity: an expanding universe of disease and genetic architecture. Nat. Rev. Genet. 25, 184–195 (2024).

5. G. Bucciol, L. Moens, B. Bosch, X. Bossuyt, J.-L. Casanova, A. Puel, I. Meyts, Lessons learned from the study of human inborn errors of innate immunity. J. Allergy Clin. Immunol. 143, 507–527 (2019).

6. A. S. Grumach, E. S. Goudouris, Inborn Errors of Immunity: how to diagnose them? J. Pediatr. (Rio J.) 97, S84–S90 (2021).

7. M. Moratti, F. Conti, M. Giannella, S. Ferrari, A. Borghesi, How to: Diagnose inborn errors of intrinsic and innate immunity to viral, bacterial, mycobacterial, and fungal infections. Clin. Microbiol. Infect. 28, 1441–1448 (2022).

8. G. Bucciol, S. Delafontaine, I. Meyts, C. Poli, Inborn errors of immunity: A field without frontiers. Immunol. Rev. 322, 15–27 (2024).

9. J. Thalhammer, G. Kindle, A. Nieters, S. Rusch, M. R. J. Seppänen, A. Fischer, B. Grimbacher, D. Edgar, M. Buckland, N. Mahlaoui, S. Ehl, K. Boztug, J. Brunner, U. F. Demel, E. Förster-Waldl, L. M. Gasteiger, L. Göschl, M. Kojić, A. Schroll, M. G. Seidel, U. Wintergerst, L. Wisgrill, S. O. Sharapova, J.-C. Goffard, T. Kerre, I. Meyts, F. Roosens, J. Smet, F. Haerynck, Z. P. Eric, V. Milenova, A. Gagro, D. Richter, Z. Chovancova, E. Hlavackova, J. Litzman, T. Milota, A. Sediva, D. A. Elaziz, R. S. Alkady, R. E. S. E. Hawary, A. S. Eldash, N. Galal, S. Lotfy, S. S. Meshaal, S. M. Reda, A. Sobh, A. Elmarsafy, M. R. J. Seppänen, P. Brosselin, V. Courteille, N. D. Vergnes, S. Kracker, M. Pergent, P. Randrianomenjanahary, G. Ahrenstorf, M. H. Albert, T. Ankermann, F. Atschekzei, U. Baumann, B. C. Becker, U. Behrends, B. H. Belohradsky, A.-K. Biegner, N. Binder, S. F. N. Bode, C. Boesecke, B. Boetticher, M. Borte, S. Borte, C. F. Classen, J. Dirks, G. Dückers, S. El-Helou, D. Ernst, M. Fasshauer, G. Fecker, K. Felgentreff, D. Foell, S. Ghosh, H. J. Girschick, S. Goldacker, N. Graf, D. Graf, J. Greil, L. G. Hanitsch, F. Hauck, M. Heeg, S. I. Heine, J. C. Henes, M. Hoenig, U. Holzer, D. Holzinger, G. Horneff, P. Hundsdoerfer, A. Jablonka, D. Jakoby, O. Joean, P. Kaiser-Labusch, C. Klemann, R. Kobbe, J. Körholz, C. M. Kramm, R. Krüger, S. Landwehr-Kenzel, K. Lehmberg, J. G. Liese, C. F. Lippert, M. E. Maccari, K. Masjosthusmann, A. Meinhardt, M. Metzler, H. Morbach, I. Müller, N. Naumann-Bartsch, J. Neubert, T. Niehues, H.-H. Peter, N. Rieber, H. Ritterbusch, J. K. Rockstroh, J. Roesler, U. Schauer, R. Scheible, M. Schmalzing, R. E. Schmidt, D. T. Schneider, S. Schreiber, C. Schuetz, A. Schulz, H. Schulze-Koops, U. Schulze-Sturm, V. Schuster, E. C. Schwaneck, K. Schwarz, C. Schwarze-Zander, M. Sirin, A. Skapenko, G. Sogkas, M. Sparber-Sauer, C. Speckmann, S. Steinmann, S. Stiehler, K. Tenbrock, H. von Bernuth, K. Warnatz, J.-C. Wasmuth, M. Weiss, T. Witte, K. Wittke, H. Wittkowski, R. A. Zeuner, E. Farmaki, M. N. Hatzistilianou, I. Kakkas, M. G. Kanariou, A. Kapousouzi, E. Liatsis, P. Maggina, E. Papadopoulou-Alataki, M. Raptaki, M. Speletas, S. Tantou, V. Goda, G. Kriván, L. Marodi, H. Abolhassani, A. Aghamohammadi, N. Rezaei, C. Feighery, T. R. Leahy, P. Ryan, N. A. Batzir, B. Z. Garty, H. Tamary, A. Aiuti, D. Amodio, C. Azzari, F. Barzaghi, L. A. Baselli, C. Cancrini, M. Carrabba, M. Cazzaniga, S. Cesaro, M. Chinello, M. G. Danieli, R. M. Dellepiane, G. Fabio, E. Gambineri, L. Lodi, V. Lougaris, C. Marasco, B. Martire, A. Marzollo, C. Milito, V. Moschese, C. Pignata, A. Plebani, F. Porta, I. Quinti, S. Ricci, A. Soresina, A. Tommasini, A. Vacca, C. Vanessa, A. Blažienė, B. Sitkauskiene, E. Gowin, E. Heropolitańska-Pliszka, B. Pietrucha, A. Szaflarska, E. Więsik-Szewczyk, B. Wolska-Kuśnierz, I. Esteves, E. Faria, L. H. Marques, J. F. Neves, S. L. Silva, C. Teixeira, S. P. da Silva, B. R. Capilna, M. N. Guseva, A. Shcherbina, A. Bobcakova, P. Ciznar, J. Gabzdilova, M. Jesenak, L. Kapustova, J. Orosova, O. Petrovicova, S. Raffac, P. Kopač, L. M. Allende, A. Antolí, G. R. Blanch, J. Carbone, R. Dieli-Crimi, M. Garcia-Prat, J. Gil-Herrera, L. I. Gonzalez-Granado, P. L. Agulló, P. Olbrich, A. Parra-Martínez, E. Paz-Artal, D. E. Pleguezuelo, N. S. Rodríguez, S. Sánchez-Ramón, J. L. Santos-Pérez, X. Solanich, P. Soler-Palacin, M. González-Amores, O. Ekwall, A. Fasth, M. Bitzenhofer-Grüber, F. Candotti, F. Dimitriou, U. Heininger, A. Holbro, P. Jandus, A. G. A. Kolios, K. Marschall, J. P. Schmid, K. M. Posfay-Barbe, S. Prader, J. Reichenbach, U. C. Steiner, J. Trück, R. G. Bredius, S. de K.-Bazen, E. de Vries, S. S. V. Henriet, T. W. Kuijpers, J. Potjewijd, A. Rutgers, K. Stol, K. J. van Aerde, J. M. V. den Berg, A. A. J. M. van de Ven, J. Montfrans, S. Aydemir, S. Baris, F. Dogu, A. Ikinciogullari, E. Karakoc-Aydiner, S. S. Kilic, A. Kiykim, Ş. İ. K. Karadağ, N. Kutukculer, S. Ocak, E. Unal, O. Boyarchuk, A. Hilfanova, L. V. Kostyuchenko, H. Alachkar, P. D. Arkwright, H. E. Baxendale, J. Bernatoniene, T. I. Coulter, T. Garcez, S. Goddard, M. M. Gompels, S. Grigoriadou, R. Herriot, A. Herwadkar, A. Huissoon, L. Ibberson, Z. Nademi, S. Noorani, S. Parvin, C. L. Steele, M. Thomas, C. Waruiru, P. F. K. Yong, H. Bourne, Initial presenting manifestations in 16,486 patients with inborn errors of immunity include infections and noninfectious manifestations. J. Allergy Clin. Immunol. 148, 1332–1341.e5 (2021).

10. A. R. Patterson, G. A. Needle, A. Sugiura, E. Q. Jennings, C. Chi, K. K. Steiner, E. L. Fisher, G. L. Robertson, C. Bodnya, J. G. Markle, R. D. Sheldon, R. G. Jones, V. Gama, J. C. Rathmell, Functional overlap of inborn errors of immunity and metabolism genes defines T cell metabolic vulnerabilities. Sci. Immunol., doi: 10.1126/sciimmunol.adh0368 (2024).

11. D. Kurup, A. M. FitzPatrick, A. Badura, I. Serra, Bridging the gap: neurodevelopmental disorder risks in inborn errors of immunity. Curr. Opin. Allergy Clin. Immunol. 24, 472–478 (2024).

12. I. Serra, M. de Koning, P. J. van der Spek, V. A. S. H. Dalm, A. Badura, Analysis of genetic overlap between inborn errors of immunity and neurodevelopmental disorders. medRxiv [Preprint] (2025). 10.1101/2025.03.01.25323148.

13. J. Wouk, D. Z. Rechenchoski, B. C. D. Rodrigues, E. V. Ribelato, L. C. Faccin-Galhardi, Viral infections and their relationship to neurological disorders. Arch. Virol. 166, 733–753 (2021).

14. N. Xiao, X. Huang, L. Chen, W. Zang, M. Guan, T. Li, I. Tuzankina, V. Chereshnev, G. Liu, Neurological Complications in Inborn Errors of Immunity: A Scoping Review of Clinical Spectrum, Pathophysiological Mechanisms, and Therapeutic Strategies. Clin. Rev. Allergy Immunol. 68, 67 (2025).

15. M. Al-Beltagi, N. K. Saeed, R. Elbeltagi, A. S. Bediwy, S. A. S. Aftab, R. Alhawamdeh, Viruses and autism: A Bi-mutual cause and effect. World J. Virol. 12, 172–192 (2023).

16. Y.-H. Hao, M. D. Fountain, K. Fon Tacer, F. Xia, W. Bi, S.-H. L. Kang, A. Patel, J. A. Rosenfeld, C. Le Caignec, B. Isidor, I. D. Krantz, S. E. Noon, J. P. Pfotenhauer, T. M. Morgan, R. Moran, R. C. Pedersen, M. S. Saenz, C. P. Schaaf, P. R. Potts, USP7 Acts as a Molecular Rheostat to Promote WASH-Dependent Endosomal Protein Recycling and Is Mutated in a Human Neurodevelopmental Disorder. Mol. Cell 59, 956–969 (2015).

17. M. C. Wimmer, H. Brennenstuhl, S. Hirsch, L. Dötsch, S. Unser, P. Caro, C. P. Schaaf, Hao-Fountain syndrome: 32 novel patients reveal new insights into the clinical spectrum. Clin. Genet. 105, 499–509 (2024).

18. F. Rafeienejad, E. Keyhani, N. Akbarfahimi, N. Nouri, Neurodevelopmental disorder: Hao– Fountain syndrome with USP7 mutation—a case report. J. Med. Case Reports 19, 363 (2025).

19. A. P. Capra, E. Agolini, M. A. La Rosa, A. Novelli, S. Briuglia, Correspondence on “Pathogenic variants in *USP7* cause a neurodevelopmental disorder with speech delays, altered behavior, and neurologic anomalies” by Fountain et al. Genet. Med. 23, 421–422 (2021).

20. M. Priolo, C. Mancini, S. Pizzi, L. Chiriatti, F. C. Radio, V. Cordeddu, L. Pintomalli, C. Mammì, B. Dallapiccola, M. Tartaglia, Complex Presentation of Hao-Fountain Syndrome Solved by Exome Sequencing Highlighting Co-Occurring Genomic Variants. Genes 13, 889 (2022).

21. M.-H. Chen, T.-P. Su, Y.-S. Chen, J.-W. Hsu, K.-L. Huang, W.-H. Chang, T.-J. Chen, Y.-M. Bai, Comorbidity of allergic and autoimmune diseases in patients with autism spectrum disorder: A nationwide population-based study. Res. Autism Spectr. Disord. 7, 205–212 (2013).

22. H. K. Hughes, R.J. Moreno, P. Ashwood, Innate immune dysfunction and neuroinflammation in autism spectrum disorder (ASD). Brain. Behav. Immun. 108, 245–254 (2023).

23. A. Hamosh, A. F. Scott, J. S. Amberger, C. A. Bocchini, V. A. McKusick, Online Mendelian Inheritance in Man (OMIM), a knowledgebase of human genes and genetic disorders. Nucleic Acids Res. 33, D514–D517 (2005).

24. A. M. Alazami, H. Hijazi, A. Y. Kentab, F. S. Alkuraya, NECAP1 loss of function leads to a severe infantile epileptic encephalopathy. J. Med. Genet. 51, 224–228 (2014).

25. E. Chouery, C. Mehawej, S. Sabbagh, J. Bleik, A. Megarbane, Early infantile epileptic encephalopathy related to NECAP1: Clinical delineation of the disease and review. Eur. J. Neurol. 29, 2486–2492 (2022).

26. V. A. Kovalskaia, V. V. Zabnenkova, M. S. Petukhova, Z. G. Markova, V. Y. Tabakov, O. P. Ryzhkova, Previously Undescribed Gross HACE1 Deletions as a Cause of Autosomal Recessive Spastic Paraplegia. Genes 13, 2186 (2022).

27. G. Sager, A. Turkyilmaz, E. A. Ates, B. Kutlubay, HACE1, GLRX5, and ELP2 gene variant cause spastic paraplegies. Acta Neurol. Belg. 122, 391–399 (2022).

28. F. Bruno, M. Anitei, D. Di Fraia, W. Durso, T. Dau, E. Cirri, M. Sannai, C. Valkova, J. Maldutyte, E. A. Miller, I. Rubio, V. Garloff, N. Kersten, G. G. Farias, A. Ori, I. Mestres, F. Calegari, C. Kaether, The microcephaly-associated protein YIPF5 differentially regulates ER export. iScience 29, 114791 (2026).

29. E. De Franco, M. Lytrivi, H. Ibrahim, H. Montaser, M. N. Wakeling, F. Fantuzzi, K. Patel, C. Demarez, Y. Cai, M. Igoillo-Esteve, C. Cosentino, V. Lithovius, H. Vihinen, E. Jokitalo, T. W. Laver, M. B. Johnson, T. Sawatani, H. Shakeri, N. Pachera, B. Haliloglu, M. N. Ozbek, E. Unal, R. Yıldırım, T. Godbole, M. Yildiz, B. Aydin, A. Bilheu, I. Suzuki, S. E. Flanagan, P. Vanderhaeghen, V. Senée, C. Julier, P. Marchetti, D. L. Eizirik, S. Ellard, J. Saarimäki-Vire, T. Otonkoski, M. Cnop, A. T. Hattersley, YIPF5 mutations cause neonatal diabetes and microcephaly through endoplasmic reticulum stress. J. Clin. Invest. 130, 6338–6353 (2020).

30. E. J. Korchak, M. Sharafi, I. Jaen Maisonet, A. Salazar-Chaparro, I. V. Semenova, H. Khan, A. L. O’Neil, P. Caro, C. P. Schaaf, S. J. Buhrlage, I. Bezsonova, Functional spectrum of USP7 pathogenic variants in Hao-Fountain syndrome: Insights into the enzyme’s activity, stability, and allosteric modulation. Proc. Natl. Acad. Sci. U. S. A. 122, e2510252122 (2025).

31. P. Lin, J. Yang, S. Wu, T. Ye, W. Zhuang, W. Wang, T. Tan, Current trends of high-risk gene Cul3 in neurodevelopmental disorders. Front. Psychiatry 14, 1215110 (2023).

32. L. van der Laan, A. Silva, L. Kleinendorst, K. Rooney, S. Haghshenas, P. Lauffer, Y. Alanay, P. Bhai, A. Brusco, S. de Munnik, B. B. A. de Vries, A. D. Vega, M. Engelen, J. C. Herkert, R. Hochstenbach, S. Hopman, S. G. Kant, R. Kira, M. Kato, B. Keren, H. Y. Kroes, M. A. Levy, N. Lock-Hock, S. M. Maas, G. M. S. Mancini, C. Marcelis, N. Matsumoto, T. Mizuguchi, A. Mussa, C. Mignot, A. Närhi, A. Nordgren, R. Pfundt, A. M. Polstra, S. Trajkova, Y. van Bever, M. José van den Boogaard, J. J. van der Smagt, T. S. Barakat, M. Alders, M. M. A. M. Mannens, B. Sadikovic, M. M. van Haelst, P. Henneman, CUL3-related neurodevelopmental disorder: Clinical phenotype of 20 new individuals and identification of a potential phenotype-associated episignature. Hum. Genet. Genomics Adv. 6, 100380 (2024).

33. S. B. Wortmann, F. M. Vaz, T. Gardeitchik, L. E. L. M. Vissers, G. H. Renkema, J. H. M. Schuurs-Hoeijmakers, W. Kulik, M. Lammens, C. Christin, L. A. J. Kluijtmans, R. J. Rodenburg, L. G. J. Nijtmans, A. Grünewald, C. Klein, J. M. Gerhold, T. Kozicz, P. M. van Hasselt, M. Harakalova, W. Kloosterman, I. Barić, E. Pronicka, S. K. Ucar, K. Naess, K. K. Singhal, Z. Krumina, C. Gilissen, H. van Bokhoven, J. A. Veltman, J. A. M. Smeitink, D. J. Lefeber, J. N. Spelbrink, R. A. Wevers, E. Morava, A. P. M. de Brouwer, Mutations in the phospholipid remodeling gene SERAC1 impair mitochondrial function and intracellular cholesterol trafficking and cause dystonia and deafness. Nat. Genet. 44, 797–802 (2012).

34. J. Finsterer, F. A. Scorza, A. C. Fiorini, C. A. Scorza, MEGDEL Syndrome. Pediatr. Neurol. 110, 25–29 (2020).

35. A. Calistri, D. Munegato, I. Carli, C. Parolin, G. Palù, The Ubiquitin-Conjugating System: Multiple Roles in Viral Replication and Infection. Cells 3, 386–417 (2014).

36. W. Jäger, S. Santag, M. Weidner-Glunde, E. Gellermann, S. Kati, M. Pietrek, A. Viejo-Borbolla, T. F. Schulz, The Ubiquitin-Specific Protease USP7 Modulates the Replication of Kaposi’s Sarcoma-Associated Herpesvirus Latent Episomal DNA. J. Virol. 86, 6745–6757 (2012).

37. S. Daubeuf, D. Singh, Y. Tan, H. Liu, H. J. Federoff, W. J. Bowers, K. Tolba, HSV ICP0 recruits USP7 to modulate TLR-mediated innate response. Blood 113, 3264–3275 (2009).

38. C. Boutell, M. Canning, A. Orr, R. D. Everett, Reciprocal Activities between Herpes Simplex Virus Type 1 Regulatory Protein ICP0, a Ubiquitin E3 Ligase, and Ubiquitin-Specific Protease USP7. J. Virol. 79, 12342–12354 (2005).

39. R. Antrobus, C. Boutell, Identification of a novel higher molecular weight isoform of USP7/HAUSP that interacts with the Herpes simplex virus type-1 immediate early protein ICP0. Virus Res. 137, 64–71 (2008).

40. D. Dutta, C. D. Williamson, N. B. Cole, J. G. Donaldson, Pitstop 2 Is a Potent Inhibitor of Clathrin-Independent Endocytosis. PLoS ONE 7, e45799 (2012).

41. J. M. Carosi, D. Denton, S. Kumar, T. J. Sargeant, Receptor Recycling by Retromer. Mol. Cell. Biol. 43, 317–334 (2023).

42. P. J. Cullen, F. Steinberg, To degrade or not to degrade: mechanisms and significance of endocytic recycling. Nat. Rev. Mol. Cell Biol. 19, 679–696 (2018).

43. F. Pontén, K. Jirström, M. Uhlen, The Human Protein Atlas--a tool for pathology. J. Pathol. 216, 387–393 (2008).

44. A. Pozhidaeva, I. Bezsonova, USP7: Structure, substrate specificity, and inhibition. DNA Repair 76, 30–39 (2019).

45. P. Rivero-Ríos, T. Tsukahara, T. Uygun, A. Chen, G. D. Chavis, S. S. P. Giridharan, S. Iwase, M. A. Sutton, L. S. Weisman, Recruitment of the SNX17-Retriever recycling pathway regulates synaptic function and plasticity. J. Cell Biol. 222, e202207025 (2023).

46. A. Singla, D. J. Boesch, H. Y. J. Fung, C. Ngoka, A. S. Enriquez, R. Song, D. A. Kramer, Y. Han, E. Banarer, A. Lemoff, P. Juneja, D. D. Billadeau, X. Bai, Z. Chen, E. E. Turer, E. Burstein, B. Chen, Structural basis for Retriever-SNX17 assembly and endosomal sorting. Nat. Commun. 15, 10193 (2024).

47. J. Zhang, Z. Jiang, A. Shi, Rab GTPases: The principal players in crafting the regulatory landscape of endosomal trafficking. Comput. Struct. Biotechnol. J. 20, 4464–4472 (2022).

48. M. D. Fernandez-Garcia, L. Meertens, M. Chazal, M. L. Hafirassou, O. Dejarnac, A. Zamborlini, P. Despres, N. Sauvonnet, F. Arenzana-Seisdedos, N. Jouvenet, A. Amara, Vaccine and Wild-Type Strains of Yellow Fever Virus Engage Distinct Entry Mechanisms and Differentially Stimulate Antiviral Immune Responses. mBio 7, e01956–01915 (2016).

49. K. M. Mayle, A. M. Le, D. T. Kamei, The Intracellular Trafficking Pathway of Transferrin. Biochim. Biophys. Acta 1820, 264–281 (2012).

50. E. M. van Dam, W. Stoorvogel, Dynamin-dependent Transferrin Receptor Recycling by Endosome-derived Clathrin-coated Vesicles. Mol. Biol. Cell 13, 169–182 (2002).

51. J. Dong, W. Tong, M. Liu, M. Liu, J. Liu, X. Jin, J. Chen, H. Jia, M. Gao, M. Wei, Y. Duan, X. Zhong, Endosomal traffic disorders: a driving force behind neurodegenerative diseases. Transl. Neurodegener. 13, 66 (2024).

52. M. Ren, G. Xu, J. Zeng, C. De Lemos-Chiarandini, M. Adesnik, D. D. Sabatini, Hydrolysis of GTP on rab11 is required for the direct delivery of transferrin from the pericentriolar recycling compartment to the cell surface but not from sorting endosomes. Proc. Natl. Acad. Sci. U. S. A. 95, 6187–6192 (1998).

53. S. Montealegre, A. Abramova, V. Manceau, A.-F. de Kanter, P. van Endert, The role of MHC class I recycling and Arf6 in cross-presentation by murine dendritic cells. Life Sci. Alliance 2, e201900464 (2019).

54. E. MacDonald, L. Brown, A. Selvais, H. Liu, T. Waring, D. Newman, J. Bithell, D. Grimes, S. Urbé, M. J. Clague, T. Zech, HRS–WASH axis governs actin-mediated endosomal recycling and cell invasion. J. Cell Biol. 217, 2549–2564 (2018).

55. K. L. Zulkefli, F. J. Houghton, P. Gosavi, P. A. Gleeson, A role for Rab11 in the homeostasis of the endosome-lysosomal pathway. Exp. Cell Res. 380, 55–68 (2019).

56. T. Bhuin, J. K. Roy, Rab11 in Disease Progression. Int. J. Mol. Cell. Med. 4, 1–8 (2015).

57. S. A. Gonçalves, T. F. Outeiro, Traffic jams and the complex role of α-Synuclein aggregation in Parkinson disease. Small GTPases 8, 78–84 (2016).

58. N. M. Gaudelli, A. C. Komor, H. A. Rees, M. S. Packer, A. H. Badran, D. I. Bryson, D. R. Liu, Programmable base editing of A•T to G•C in genomic DNA without DNA cleavage. Nature 551, 464–471 (2017).

59. R. T. Walton, K. A. Christie, M. N. Whittaker, B. P. Kleinstiver, Unconstrained genome targeting with near-PAMless engineered CRISPR-Cas9 variants. Science 368, 290–296 (2020).

60. M. G. Kluesner, D. A. Nedveck, W. S. Lahr, J. R. Garbe, J. E. Abrahante, B. R. Webber, B. S. Moriarity, EditR: A Method to Quantify Base Editing from Sanger Sequencing. CRISPR J. 1, 239–250 (2018).

61. R. Hamel, O. Dejarnac, S. Wichit, P. Ekchariyawat, A. Neyret, N. Luplertlop, M. Perera-Lecoin, P. Surasombatpattana, L. Talignani, F. Thomas, V.-M. Cao-Lormeau, V. Choumet, L. Briant, P. Desprès, A. Amara, H. Yssel, D. Missé, Biology of Zika Virus Infection in Human Skin Cells. J. Virol. 89, 8880–8896 (2015).

62. D. Olagnier, M. Muscolini, C. B. Coyne, M. S. Diamond, J. Hiscott, Mechanisms of Zika Virus Infection and Neuropathogenesis. DNA Cell Biol. 35, 367–372 (2016).

63. M. D. Fountain, D. S. Oleson, M. E. Rech, L. Segebrecht, J. V. Hunter, J. M. McCarthy, P. J. Lupo, M. Holtgrewe, R. Moran, J. A. Rosenfeld, B. Isidor, C. Le Caignec, M. S. Saenz, R. C. Pedersen, T. M. Morgan, J. P. Pfotenhauer, F. Xia, W. Bi, S.-H. L. Kang, A. Patel, I. D. Krantz, S. E. Raible, W. Smith, I. Cristian, E. Torti, J. Juusola, F. Millan, I. M. Wentzensen, R. E. Person, S. Küry, S. Bézieau, K. Uguen, C. Férec, A. Munnich, M. van Haelst, K. D. Lichtenbelt, K. van Gassen, T. Hagelstrom, A. Chawla, D. L. Perry, R. J. Taft, M. Jones, D. Masser-Frye, D. Dyment, S. Venkateswaran, C. Li, L. F. Escobar, D. Horn, R. C. Spillmann, L. Peña, J. Wierzba, T. M. Strom, I. Parenti, F. J. Kaiser, N. Ehmke, C. P. Schaaf, Pathogenic variants in USP7 cause a neurodevelopmental disorder with speech delays, altered behavior, and neurologic anomalies. Genet. Med. 21, 1797–1807 (2019).

64. Y.-H. Chan, Z. Liu, P. Bastard, N. Khobrekar, K. M. Hutchison, Y. Yamazaki, Q. Fan, D. Matuozzo, O. Harschnitz, N. Kerrouche, K. Nakajima, P. Amin, A. Yatim, D. Rinchai, J. Chen, P. Zhang, G. Ciceri, J. Chen, K. Dobbs, S. Belkaya, D. Lee, A. Gervais, K. Aydın, A. Kartal, M. L. Hasek, S. Zhao, E. G. Reino, Y. S. Lee, Y. Seeleuthner, M. Chaldebas, R. Bailey, C. Vanhulle, L. Lorenzo, S. Boucherit, F. Rozenberg, N. Marr, T. H. Mogensen, M. Aubart, A. Cobat, O. Dulac, M. Emiroglu, S. R. Paludan, L. Abel, L. Notarangelo, R. Longnecker, G. Smith, L. Studer, J.-L. Casanova, S.-Y. Zhang, Human TMEFF1 is a restriction factor for herpes simplex virus in the brain. Nature 632, 390–400 (2024).

65. Y. Dai, M. Idorn, M. C. Serrero, X. Pan, E. A. Thomsen, R. Narita, M. Maimaitili, X. Qian, M. B. Iversen, L. S. Reinert, R. K. Flygaard, M. Chen, X. Ding, B. Zhang, M. E. Carter-Timofte, Q. Lu, Z. Jiang, Y. Zhong, S. Zhang, L. Da, J. Zhu, M. Denham, P. Nissen, T. H. Mogensen, J. G. Mikkelsen, S.-Y. Zhang, J.-L. Casanova, Y. Cai, S. R. Paludan, TMEFF1 is a neuron-specific restriction factor for herpes simplex virus. Nature 632, 383–389 (2024).

66. J. G. Doench, E. Hartenian, D. B. Graham, Z. Tothova, M. Hegde, I. Smith, M. Sullender, B. L. Ebert, R. J. Xavier, D. E. Root, Rational design of highly active sgRNAs for CRISPR-Cas9-mediated gene inactivation. Nat. Biotechnol. 32, 1262–1267 (2014).

67. 67. R. E. Hanna, M. Hegde, C. R. Fagre, P. C. DeWeirdt, A. K. Sangree, Z. Szegletes, A. Griffith, M. N. Feeley, K. R. Sanson, Y. Baidi, L. W. Koblan, D. R. Liu, J. T. Neal, J. G. Doench, Massively parallel assessment of human variants with base editor screens. Cell 184, 1064–1080.e20 (2021).

68. A. K. Sangree, A. L. Griffith, Z. M. Szegletes, P. Roy, P. C. DeWeirdt, M. Hegde, A. V. McGee, R. E. Hanna, J. G. Doench, Benchmarking of SpCas9 variants enables deeper base editor screens of BRCA1 and BCL2. Nat. Commun. 13, 1318 (2022).

69. T. Doyle, O. Moncorgé, B. Bonaventure, D. Pollpeter, M. Lussignol, M. Tauziet, L. Apolonia, M.-T. Catanese, C. Goujon, M. H. Malim, The interferon-inducible isoform of NCOA7 inhibits endosome-mediated viral entry. Nat. Microbiol. 3, 1369–1376 (2018).

70. M. C. Schwarz, M. Sourisseau, M. M. Espino, E. S. Gray, M. T. Chambers, D. Tortorella, M. J. Evans, Rescue of the 1947 Zika Virus Prototype Strain with a Cytomegalovirus Promoter-Driven cDNA Clone. mSphere 1, e00246–16 (2016).

