## Supplementary figures for "The Hao-Fountain syndrome gene USP7 restricts neurotropic orthoflavivirus entry through cell-intrinsic control of endosomal dynamics"

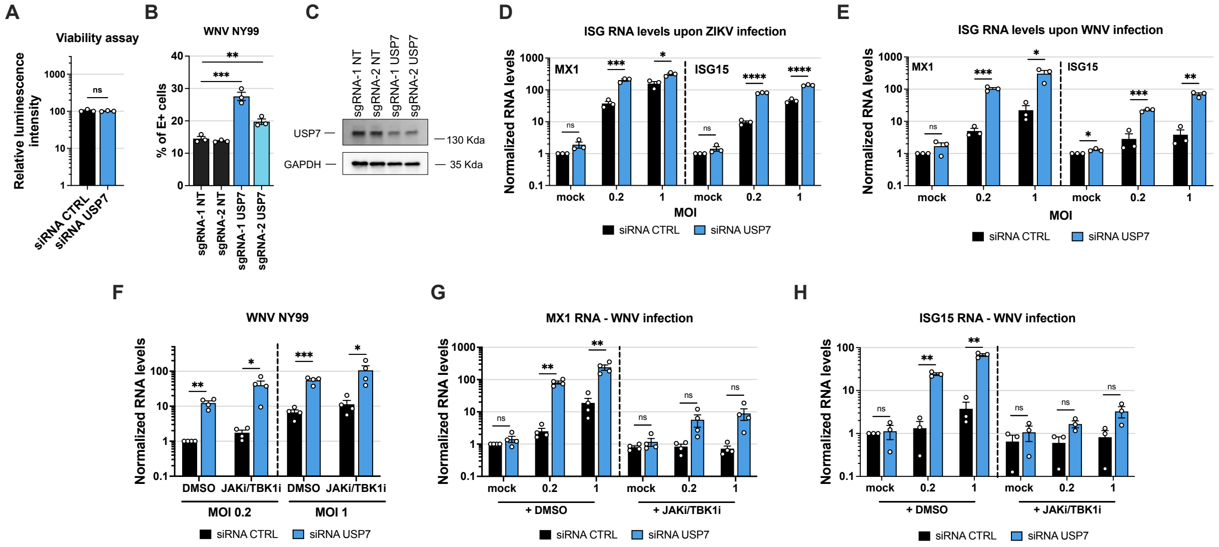


**Fig. S1. USP7 antiviral activity is independent on the type-I interferon response. (A)** Cell viability assays in siRNA-transfected A549 cells used in figure 2. **(B)** West Nile virus (WNV) strain NY99 infection rates at 24 h post infection were quantified in A549-Cas9 cells expressing non-targeting sgRNAs (sgRNA-1 NT, sgRNA-2 NT) or sgRNAs targeting USP7 (sgRNA-1 USP7, sgRNA-2 USP7). Infection was measured by intracellular staining of the viral Envelop (E) protein and flow cytometry analysis of cells infected at a multiplicity of infection (MOI) of 0.1. Multiple unpaired t-tests with false discovery rate correction were used for statistical analysis. **(C)** USP7 silencing efficiency, measured by immunoblot using corresponding samples from panel B. **(D)** Relative quantification of *MX1* and *ISG15* mRNA levels by RT-qPCR (normalized to GAPDH) in control (siRNA CTRL) or USP7-depleted (siRNA USP7) A549 cells infected with Zika virus (ZIKV) strain MR-766 for 24 h at MOI 0.2 or 1. Multiple unpaired t-tests with false discovery rate correction were used for statistical analysis. **(E)** Relative quantification of *MX1* and *ISG15* mRNA levels by RT-qPCR (normalized to GAPDH) in siRNA CTRL or siRNA USP7 A549 cells infected with WNV strain NY99 for 24 h at MOI 0.2 or 1. Multiple unpaired t-tests with false discovery rate correction were used for statistical analysis. **(F)** WNV strain NY99 infection rates at 24 h post infection in A549 cells transfected with siRNA CTRL or siRNA USP7 and pre-treated for 1 h with 500 nM

TBK1 and pan-JAK inhibitors (+JAKi/TBK1i) or an equivalent volume of DMSO. Cells were infected at the indicated MOI, and infection was measured by RT-qPCR. Multiple unpaired t-tests with false discovery rate correction were used for statistical analysis. **(G)** Relative quantification of *MX1* mRNA levels by RT-qPCR (normalized to GAPDH) in siRNA CTRL o siRNA USP7) A549 cells pre-treated 1 h with 500 nM TBK1 and pan-JAK inhibitors (+JAKi/TBK1i) or an equivalent volume of DMSO and infected with WNV strain NY99 for 24 h at indicated MOI. Multiple unpaired t-tests with false discovery rate correction were used for statistical analysis. **(H)** Relative quantification of *ISG15* mRNA levels by RT-qPCR (normalized to GAPDH) in siRNA CTRL or siRNA USP7 A549 cells pre-treated 1 h with 500 nM TBK1 and pan-JAK inhibitors (+JAKi/TBK1i) or an equivalent volume of DMSO and infected with WNV strain NY99 for 24 h at indicated MOI. Multiple unpaired t-test with false discovery rate correction were used for statistical analysis. Data information: (A-B, D-H) Mean ± SEM of three or four biological replicates. p-values are denoted as follows: ns, not significant, *p < 0.05, **p < 0.01, ***p < 0.001, ****p < 0.0001.

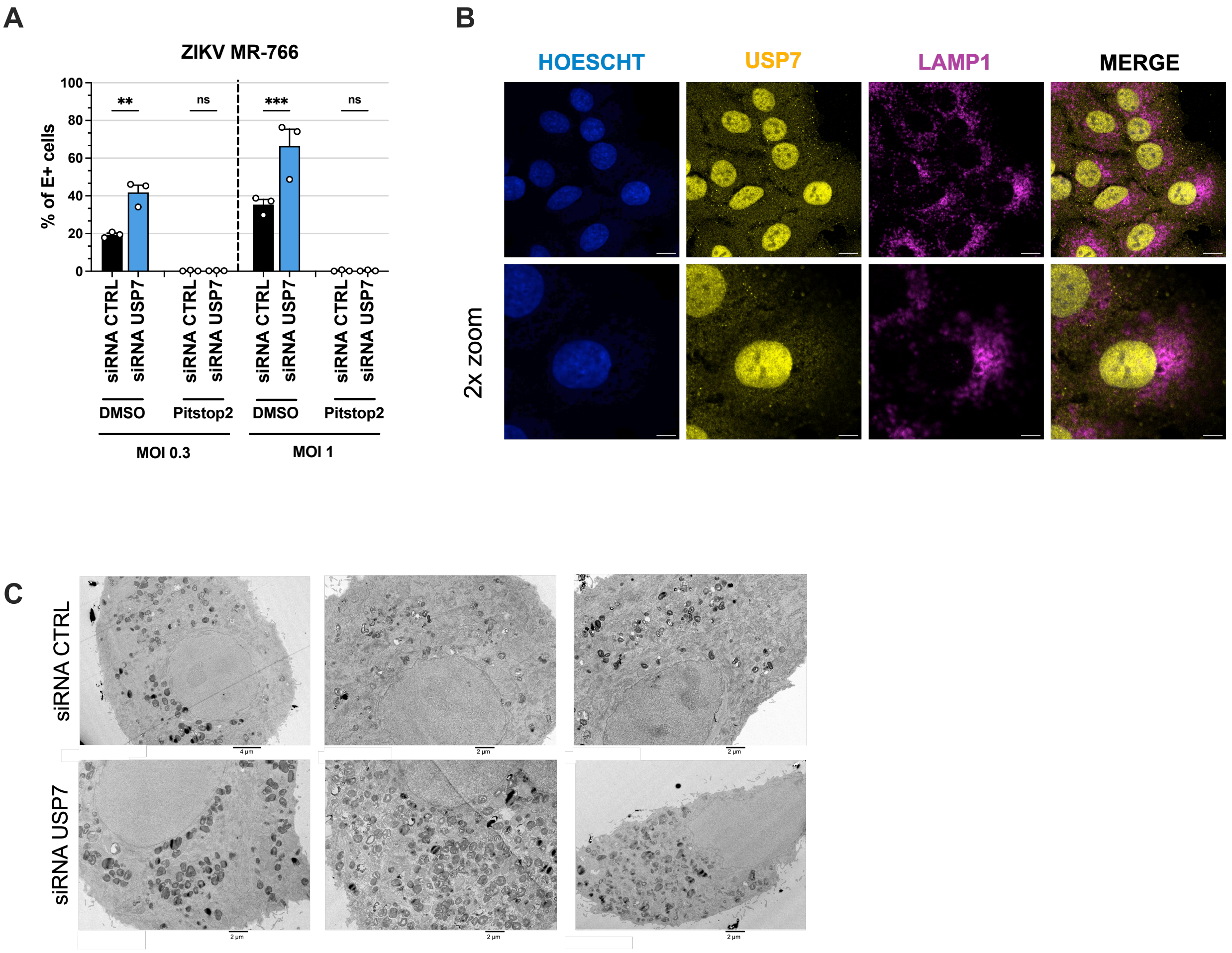


**Fig. S2. USP7 silencing does not impact ZIKV entry route. (A)** West Nile virus (WNV) strain NY99 infection at 24 h post-infection in A549 cells transfected with siRNA CTRL or siRNA USP7 and pre-treated with the indicated concentrations of Pitstop2 or equivalent volumes of DMSO as vehicle control. Infection was quantified by intracellular staining of viral E protein and flow cytometry in cells infected at an MOI of 0.1. Statistical analysis was performed using multiple unpaired t-tests with false discovery rate correction. Mean ± SEM of three biological replicates **(B)** Confocal microscopy of A549 cells stained with anti-LAMP1 (purple) and anti-USP7 (yellow) antibodies, and Hoescht for nuclei. Top: scale bar: 10 µm. Bottom: scale bar: 20 µm. **(C)** Electron micrographs of siRNA CTRL and siRNA USP7 A549 cells. Scale bars: 2 or 4 µm, as indicated in the images.

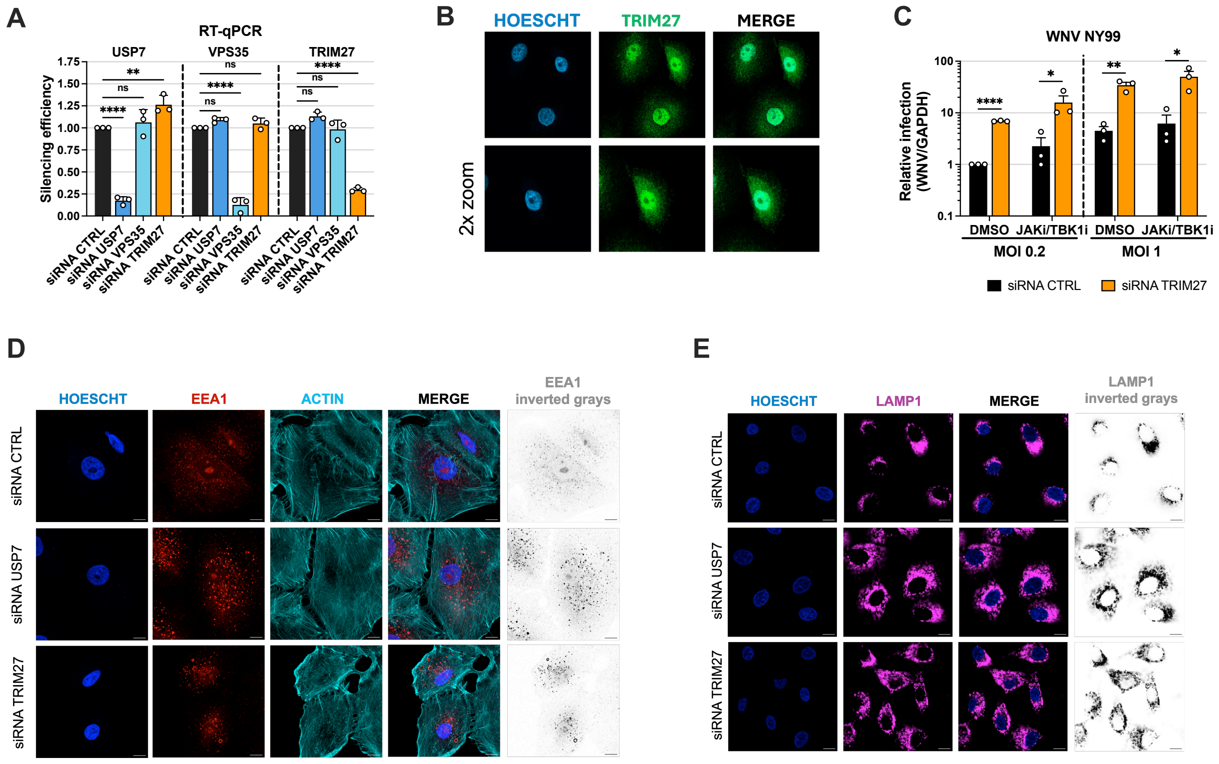


**Fig. S3. TRIM27 silencing efficiency and effects of TRIM27 depletion on early and late endosomal compartments. (A)** USP7, TRIM27, and VPS35 silencing efficiency in A549 cells transfected with siRNA pools, measured by RT-qPCR. Two-way ANOVA with Tukey's test. **(B)** Confocal microscopy of A549 cells stained with anti-TRIM27 (green) and Hoescht for nuclei. Top: scale bar: 10 µm. Bottom: scale bar: 20 µm. **(C)** WNV strain NY99 infection rates at 24 h post infection in A549 cells transfected with siRNA CTRL or siRNA TRIM27 and pre-treated for 1 h with 500 nM TBK1 and pan-JAK inhibitors (+JAKi/TBK1i) or an equivalent volume of DMSO. Cells were infected at the indicated MOIs, and infection was measured by RT-qPCR. Multiple unpaired t-tests with false discovery rate correction. **(D)** Confocal microscopy of siRNA CTRL, siRNA USP7 and siRNA TRIM27 A549 cells stained with anti-EEA1 (red and inverted gray), phalloidin for F-actin (cyan), and Hoescht for nuclei (blue). Scale bar: 10 µm. **(E)** Confocal microscopy of siRNA CTRL, siRNA USP7 and siRNA TRIM27 A549 cells stained with anti-LAMP1 (purple and inverted gray) and Hoescht for nuclei (blue). Scale bar: 10 µm.

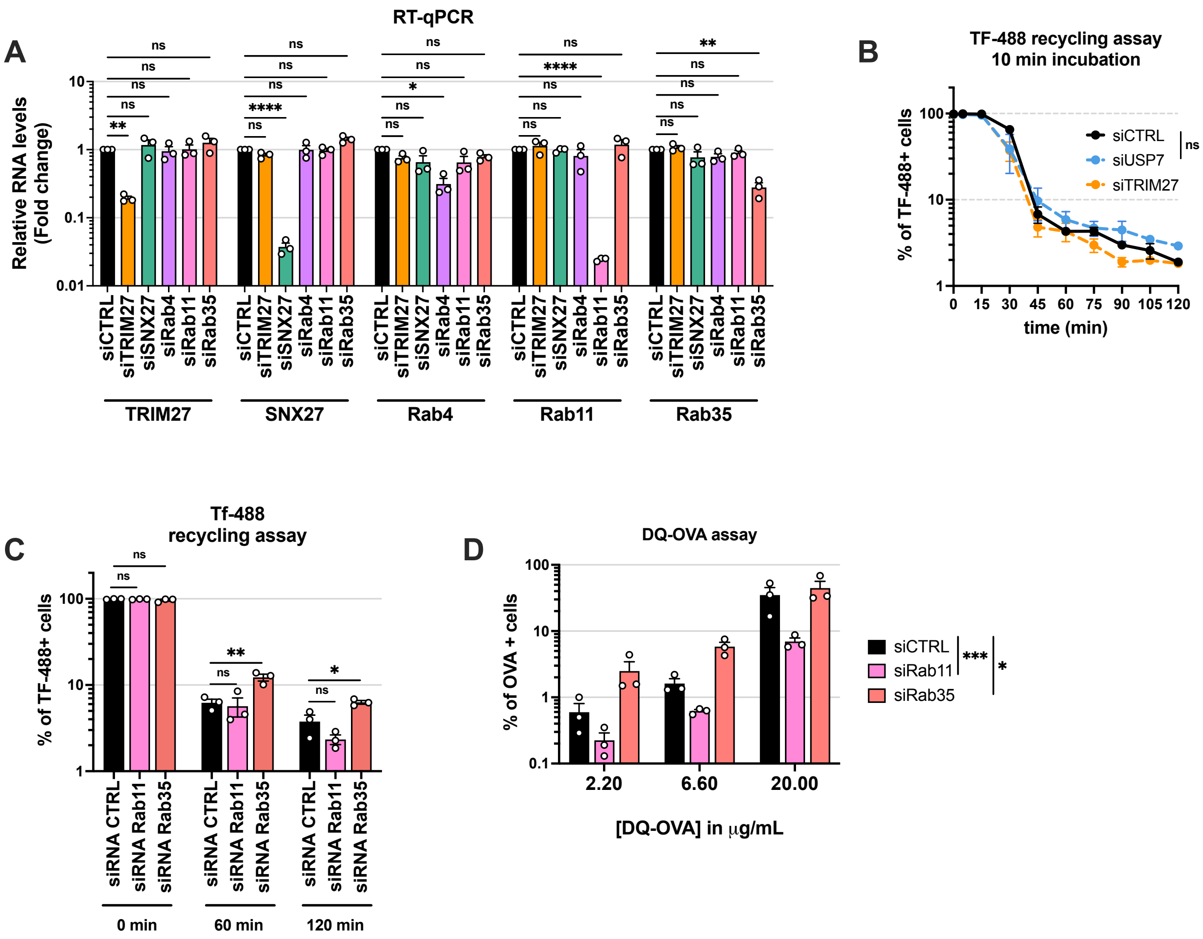


**Figure S4. USP7 and TRIM27 depletion in A549 cells do not impair transferrin recycling at early time points. (A)** TRIM27, SNX27, Rab4A, Rab11A and Rab35/a silencing efficiency in A549 cells transfected with siRNA pools, measured by RT-qPCR. Two-way ANOVA with Tukey's test. **(B)** Tf-488 recycling assays in A549 cells transfected with siRNA CTRL, siRNA USP7, or siRNA TRIM27. Cells were incubated with 15 µg/mL Tf-488 for 30 min at 4 °C, shifted to 37 °C for 10 min, washed, and incubated with unlabeled transferrin before fixation at the indicated time points to quantify the percentage of Tf-488$^{+}$ cells by flow cytometry. Statistical analysis was performed using simple linear regression analysis. **(C)** Tf-488 recycling assays in A549 cells transfected with siRNA CTRL, siRNA Rab11A, or siRNA Rab35A. Cells were incubated with 25 µg/mL Tf-488 for 30 min at 4 °C, shifted to 37 °C for 1 h, washed, and incubated with unlabeled transferrin before fixation at the indicated time points to quantify the percentage of Tf-488$^{+}$ cells by flow cytometry. Multiple unpaired t-tests with false discovery rate correction. **(D)** DQ-OVA assays in A549 cells transfected with siRNA CTRL, siRNA USP7, or siRNA TRIM27. Cells were incubated with the indicated concentrations of DQ-OVA for 15 min at 37 °C, washed, and fixed to quantify the percentage of cells positive for DQ-OVA fluorescence (degraded OVA) by flow cytometry. Simple linear regression analysis on log-transformed data. Data information: **(A-D)** Mean ± SEM of three biological replicates. p-values are denoted as follows: ns, not significant, *p < 0.05, **p < 0.01, ***p < 0.001, ****p < 0.0001.

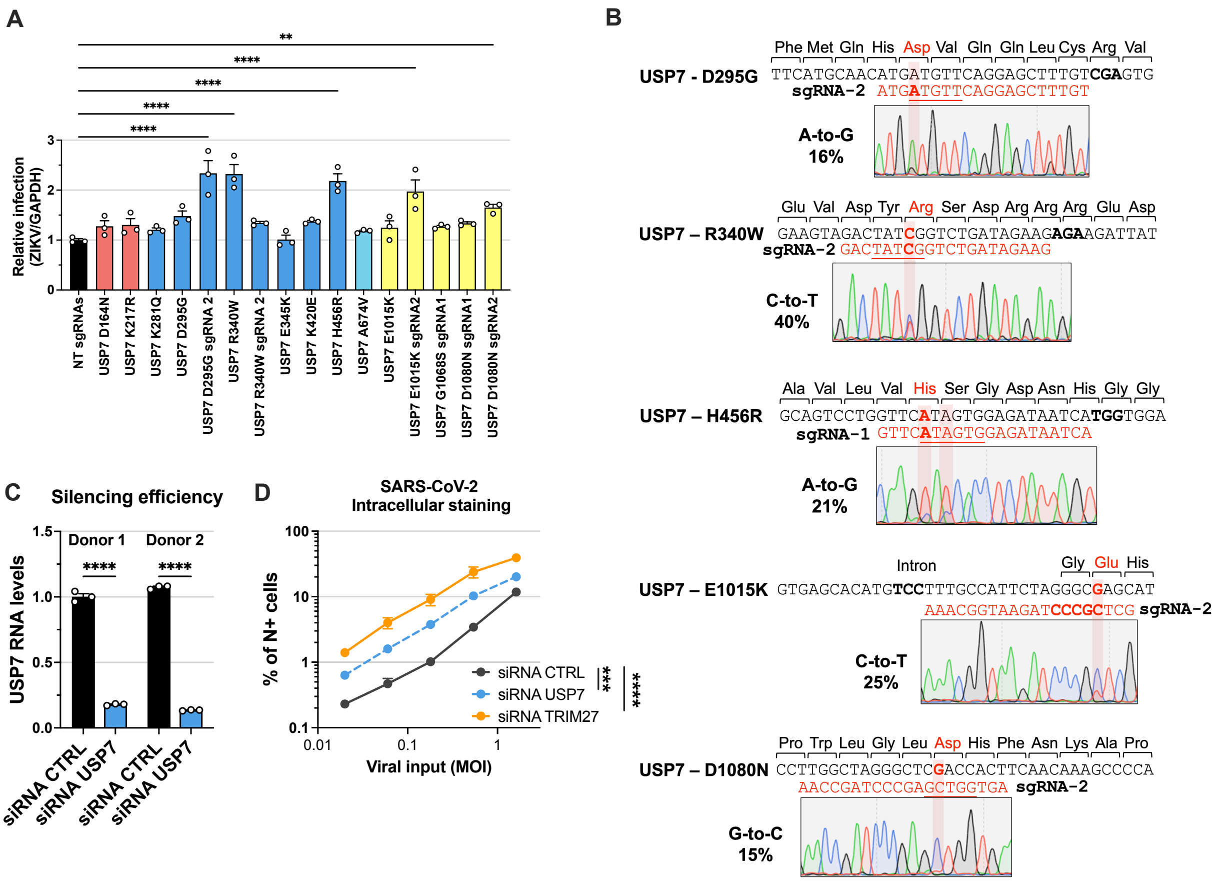


**Fig. S5. CRISPR/Cas9 base-editing efficiencies, silencing efficiencies in primary fibroblasts and SARS-CoV-2 infection. (A)** Zika virus (ZIKV) strain MR-766 infection at 24 h post-infection in A549 cells expressing Cas9 base editors and sgRNAs introducing the indicated mutations. Infection was measured by RT-qPCR and normalized to GAPDH mRNA levels. Statistical analysis was performed using one-way ANOVA with Dunnett's test. **(B)** Sequencing analysis of PCR amplicons from genomic DNA of A549 cells edited to introduce the indicated mutations (D295G, R340W, H456R, E1015K, and D1080N). The target locus is shown in black, with the target amino acid-coding sequence in red and the PAM sequence in bold. The homologous region of the sgRNAs is indicated in red, with the target nucleotide in bold and the Cas9 editor activity window underlined. Editing efficiency, expressed as a percentage, was quantified from peak overlaps using the EditR web tool. **(C)** USP7 silencing efficiency in human primary fibroblasts from two donors measured by RT-qPCR. Two-way ANOVA with Tukey's test. **(D)** SARS-CoV-2 strain WA1 infection rates at 24 h post infection in A549-Ace2 cells transfected with siRNA CTRL, siRNA USP7 or siRNA TRIM27. Infection was measured by intracellular staining of the viral N protein and flow cytometry analysis of cells infected at indicated MOI. Multiple unpaired t-tests with false discovery rate correction were used for statistical analysis. Data information: (A, C and D) Mean ± SEM of three biological replicates. p-values are denoted as follows: ns, not significant, $^*p < 0.05$, $^{**}p < 0.01$, $^{***}p < 0.001$, $^{****}p < 0.0001$.
