## Supplementary tables for "The Hao-Fountain syndrome gene USP7 restricts neurotropic orthoflavivirus entry through cell-intrinsic control of endosomal dynamics"

### SUPPLEMENTAL TABLE 1

*Screen targets and diseases*

| Symbol | Function | Disease | OMIM ID |
|---|---|---|---|
| STAMBP | Deubiquitinating isopeptidase | Microcephaly-capillary malformation syndrome | 606247 |
| LMBRD1 | Lysosomal exporter | Methylmalonic aciduria and homocystinuria | 612625 |
| VPS50 | Endosome-associated recycling protein (EARP) complex | Neurodevelopmental disorder with microcephaly, seizures, and neonatal cholestasis | 616465 |
| NECAP1 | Clathrin-mediated endocytosis | Developmental and epileptic encephalopathy | 611623 |
| TBC1D2B | GTPase-activating protein | Neurodevelopmental disorder with seizures and gingival overgrowth | 619152 |
| CC2D1A | Endocytosis regulator | Intellectual developmental disorder, autosomal recessive | 610055 |
| VAMP2 | Vesicle trafficking / membrane fusion | Neurodevelopmental disorder with hypotonia and autistic features | 185881 |
| REEP1 | Endoplasmic reticulum morphology | Neuronopathy, distal hereditary motor, autosomal recessive | 609139 |
| BAP1 | Ubiquitin carboxy-terminal hydrolase | Kury-Isidor syndrome | 603089 |
| COPB2 | Subunit of the Golgi coatomer complex | Microcephaly, primary, autosomal recessive | 606990 |
| YIPF5 | COPII-dependent trafficking | Microcephaly, epilepsy, and diabetes syndrome | 611483 |
| HACE1 | HECT domain-containing E3 ubiquitin ligase | Spastic paraplegia | 610876 |
| SERAC1 | Intracellular cholesterol trafficking | Encephalopathy | 614725 |
| TMTC3 | Endoplasmic reticulum stress response | Lissencephaly | 617218 |
| TRAPPC6B | Vesicle transport | Neurodevelopmental disorder with microcephaly, epilepsy, and brain atrophy | 610397 |
| IER3IP1 | Endoplasmic reticulum stress response | Microcephaly, epilepsy, and diabetes syndrome | 609382 |
| VPS13B | Golgi apparatus | Cohen syndrome | 607817 |
| RIC1 | Rab6a activator | CATIFA syndrome | 610354 |
| INPP5E | Golgi phosphatase | Joubert syndrome 1 | 613037 |
| MAN1B1 | ER-Golgi Mannosidase | Rafiq syndrome | 604346 |
| TMEM251 | Lysosomal enzyme transport | Dysostosis multiplex, Ain-Naz type | 619332 |
| HECW2 | HECT-type ubiquitin ligase | Neurodevelopmental disorder with hypotonia | 617245 |
| CUL3 | Scaffolding of the Cullin-RING ligase complex | Neurodevelopmental disorder with or without autism or seizures | 603136 |
| USP7 | Ubiquitin hydrolase | Hao-Fountain syndrome | 602519 |

#### SUPPLEMENTAL TABLE 2

*Oligonucleotides used for molecular cloning*

| Gene | Oligo | Sequence |
|---|---|---|
| Cas9-NG | PCR_fw | CCATTTCAGGTGTCGTGAGGATGAAATCTTCTCACCATCACCATCACCATG |
| | PCR_rev | GAGGTTGATTGTCGACTTAAGGTTAGGCATCAGCAAACCCAAG |
| Non-targeting | Top oligo sgRNA 1 | CACCGCCCCGCCGCCCTCCCCTCC |
| | Bottom oligo sgRNA 1 | AAACGGAGGGGAGGGCGGCGGGGC |
| | Top oligo sgRNA 2 | CACCGTATTACTGATATTGGTGGG |
| | Bottom oligo sgRNA 2 | AAACCCCACCAATATCAGTAATAC |
| USP7 | Top oligo sgRNA 1 | CACCGACCATACCCAAATTATTCCG |
| | Bottom oligo sgRNA 1 | AAACCGGAATAATTTGGGTATGGTC |

#### SUPPLEMENTAL TABLE 3

*Oligonucleotides used for CRISPR/Cas9 base editing*

| Gene | Vector | Oligo | Sequence |
|---|---|---|---|
| Non-targeting control | pRDA_429-CTRL guide 1 | Top | CACCGCCCCGCCGCCCTCCCCTCC |
| | | Bottom | AAACGGAGGGGAGGGCGGCGGGGC |
| | pRDA_429-CTRL guide 2 | Top | CACCGTATTACTGATATTGGTGGG |
| | | Bottom | AAACCCCACCAATATCAGTAATAC |
| | pRDA_78 CBE4-Cas9-NG-CTRL guide 1 | Top | CACCGCCCCGCCGCCCTCCCCTCC |
| | | Bottom | AAACGGAGGGGAGGGCGGCGGGGC |
| | pRDA_78 CBE4-Cas9-NG-CTRL guide 2 | Top | CACCGTATTACTGATATTGGTGGG |
| | | Bottom | AAACCCCACCAATATCAGTAATAC |
| USP7 | pRDA_78_CBE4-Cas9-NG-USP7 D164N | Top | CACCCCAATCATTTTCTTTATGGA |
| | | Bottom | AAACTCCATAAAGAAAATGATTGG |
| | pRDA_429 ABE8e-Cas9-NG-USP7 K217R | Top | CACCTAAAGAATCAGGGAGCGACT |
| | | Bottom | AAACAGTCGCTCCCTGATTCTTTA |
| | pRDA_429 ABE8e-Cas9-NG-USP7 K281Q | Top | CACCTTAACAAAGTCATTTGGGTA |
| | | Bottom | AAACTACCCAAATGACTTTGTTAA |
| | pRDA_429 ABE8e-Cas9-NG-USP7 D295G guide 1 | Top | CACCAACATGATGTTCAGGAGCTT |
| | | Bottom | AAACAAGCTCCTGAACATCATGTT |
| | pRDA_429 ABE8e-Cas9-NG-USP7 D295G guide 2 | Top | CACCATGATGTTCAGGAGCTTTGT |
| | | Bottom | AAACACAAAGCTCCTGAACATCAT |
| | pRDA_78_CBE4-Cas9-NG-USP7 R340W guide 1 | Top | CACCGACTATCGGTCTGATAGAAG |
| | | Bottom | AAACCTTCTATCAGACCGATAGTC |
| | pRDA78_CBE4-Cas9-NG-USP7 R340W guide 2 | Top | CACCTATCGGTCTGATAGAAGAGA |
| | | Bottom | AAACTCTCTTCTATCAGACCGATA |
| | pRDA_78_CBE4-Cas9-NG-USP7 E345K | Top | CACCAATCTTCTCTTCTATCAGAC |
| | | Bottom | AAACGTCTGATAGAAGAGAAGATT |
| | pRDA_429 ABE8e-Cas9-NG-USP7 K420E | Top | CACCAAATATCAAGATCAATGATA |
| | | Bottom | AAACTATCATTGATCTTGATATTT |
| | pRDA_429 ABE8e-Cas9-NG-USP7 H456R | Top | CACCGTTCATAGTGGAGATAATCA |
| | | Bottom | AAACTGATTATCTCCACTATGAAC |
| | pRDA_78_CBE4-Cas9-NG-USP7 A674V | Top | CACCGGAGCGACCTTACCCAAGTT |
| | | Bottom | AAACAACTTGGGTAAGGTCGCTCC |
| | pRDA78_CBE4-Cas9-NG-USP7 E1015K guide 1 | Top | CACCGCTCGCCCTAGAATGGCAAA |
| | | Bottom | AAACTTTGCCATTCTAGGGCGAGC |
| | pRDA78_CBE4-Cas9-NG-USP7 E1015K guide 2 | Top | CACCTGCTCGCCCTAGAATGGCAA |
| | | Bottom | AAACTTGCCATTCTAGGGCGAGCA |
| | pRDA78_CBE4-Cas9-NG-USP7 G1068S | Top | CACCCTTACCGGGCTGTGGCTCAA |
| | | Bottom | AAACTTGAGCCACAGCCCGGTAAG |
| | pRDA_78_CBE4-Cas9-NG-USP7 D1080N guide 1 | Top | CACCAGTGGTCGAGCCCTAGCCAA |
| | | Bottom | AAACTTGGCTAGGGCTCGACCACT |
| | pRDA_78_CBE4-Cas9-NG-USP7 D1080N guide 2 | Top | CACCAAGTGGTCGAGCCCTAGCCA |
| | | Bottom | AAACTGGCTAGGGCTCGACCACTT |

**SUPPLEMENTAL TABLE 4**

*Oligonucleotides used for genomic DNA PCR and sequencing*

| Mutation | Oligo | Sequence |
|---|---|---|
| USP7 D164N | PCR_fw | GTCTTGCCATGCACAAGCAG |
| | PCR_rev | GTGAGACGTGAATCCAGAAAGC |
| | Sequencing | GGGAACAACAAGCAGTAATGCACC |
| USP7 K217R | PCR_fw | ACACACACTGCAGGAGCAAT |
| | PCR_rev | GTTGGCGGTGGTAACTCTGA |
| | Sequencing | CCCCTCAACCTTGAACTGATCTTTC |
| USP7 K281Q | PCR_fw | GAGACTTCGTGTCACGTGGG |
| | PCR_rev | GGACAGCCAACAAAGCCAGTC |
| | Sequencing | GCTGTGTACATGATGCCAACCG |
| USP7 D295G | PCR_fw | GAGACTTCGTGTCACGTGGG |
| | PCR_rev | GGACAGCCAACAAAGCCAGTC |
| | Sequencing | CCCTATTCAAGTAATTCTTACCAGCC |
| USP7 R340W and E345K | PCR_fw | TGGTGTGGACCATTACGAGTT |
| | PCR_rev | CTGGAGTTCGAGGCTGCACTG |
| | Sequencing | GCACCTGTGTAGAGGGCACC |
| USP7 K420E | PCR_fw | CTTCCCTGATTGCTGGGCATC |
| | PCR_rev | AGCACTGAGACTAACCCCCT |
| | Sequencing | GTTTAGGAAGCAGAGAAAGGTGTGA |
| USP7 H456R | PCR_fw | CCCAGAGCAGTTACCACTTGA |
| | PCR_rev | GAAACAAGGTAGGACCAGGGG |
| | Sequencing | GGGTCCCACCACTTACTTTG |
| USP7 A674V | PCR_fw | CGTGCAGAAGGCGTTAGTCCTC |
| | PCR_rev | GGAAACTCGCAGCCAAGTCAG |
| | Sequencing | CAGGAATCCAACGCTACTGCTC |
| USP7 E1015K | PCR_fw | CTTGTCACAGTGGCGCATTTCC |
| | PCR_rev | TAACACCAGCAGCGAATCCTC |
| | Sequencing | CCCAGCTGCACACCTTCTC |
| USP7 G1068S | PCR_fw | CTTGTCACAGTGGCGCATTTCC |
| | PCR_rev | TAACACCAGCAGCGAATCCTC |
| | Sequencing | GCAATTGTAATGATGGGCCGAC |
| USP7 D1080N | PCR_fw | CTTGTCACAGTGGCGCATTTCC |
| | PCR_rev | TAACACCAGCAGCGAATCCTC |
| | Sequencing | CCTTGAACACACCAGCTTGGAAATC |

**SUPPLEMENTAL TABLE 5**

*Oligonucleotides used for RT-qPCR*

| Target | Oligo | Sequence |
|---|---|---|
| GAPDH | Forward | ATGTTCCAATATGATTCCACCC |
| | Reverse | ATCGCCCCACTTGATTTTG |
| WNV_NY99 | Forward | GAGTTGATGTGCGGCTTGAT |
| | Reverse | GCACTAATCGCGAGACAGAC |
| ZIKV_MR-766 | Forward | AAATACACATACCAAAACAAAGTGGT |
| | Reverse | TCCRCTCCCYCTYTGGTCTTG |
| YFV_17D | Forward | AGGTCCAGTTGATCGCGGC |
| | Reverse | GAGCGACAGCCCCGATTTCT |
| DENV_2 | Forward | TCCATGCAAGATCCCTTTTGA |
| | Reverse | ATGGCCATTCTCTTCGCCCC |
| MX1 | Forward | TCCAGCCACATCCCTTTGAT |
| | Reverse | TCCTTCAGGAACTTCCGCTT |
| ISG15 | Forward | ACAAATGCGACGAACCTCTG |
| | Reverse | AAGGTCAGCCAGAACAGGTC |
| IFNB | Forward | GTCTCCTCCAAATTGCTCTC |
| | Reverse | ACAGGAGCTTCTGACACTGA |
| IL6 | Forward | GGCACTGGCAGAAAACAACC |
| | Reverse | GCAAGTCTCCTCATTGAATCC |
| USP7 | Forward | GGAGGAGGACATGGAGGATG |
| | Reverse | CTTCCATGGCAGATTTCGCA |
| TRIM27 | Forward | ACCTCCCCAATGACTGCCCT |
| | Reverse | CGGACAGGCCCACAAAAGGT |
| VPS35 | Forward | CCGTGGTGTGCAACATCCCT |
| | Reverse | TGATGCTGCATTCGCACCCA |
| SNX27 | Forward | GAGCAGGCGAGAAGGAATTG |
| | Reverse | GCTTAGAACACAGCTGCCTC |
| RAB4A | Forward | ACTAGCACTAGGGATTCTGG |
| | Reverse | AGAATGTGTTTTCTAGCAGG |
| RAB11A | Forward | ACATCAGCATATTATCGTGG |
| | Reverse | GACGTAGATCACTCTTATTGC |
| RAB35A | Forward | TGTCAACGTCAAGCGATGG |
| | Reverse | GGTCATCATTCTTATTGCCCACT |

**SUPPLEMENTAL TABLE 6**
*siRNA targets and ID - Horizon Dharmacon*

| Target | Product | Gene Accession | Catalog ID |
|---|---|---|---|
| CTRL1 | siGENOME Non-targeting Control SMARTPool | | D-001206-13 |
| CTRL2 | siGENOME Non-targeting Control SMARTPool | | D-001206-14 |
| COPB2 | siGENOME Human COPB2 SMARTPool | NM_004766 | M-005791-00 |
| TBC1D2B | siGENOME Human TBC1D2B SMARTPool | NM_015079 | M-014127-00 |
| VAPB | siGENOME Human VAPB SMARTPool | NM_004738 | M-017795-00 |
| TRAPPC6B | siGENOME Human TRAPPC6B SMARTPool | NM_177452 | M-018297-01 |
| TMTC3 | siGENOME Human TMTC3 SMARTPool | NM_181783 | M-018618-01 |
| C15orf57 | siGENOME Human C15orf57 SMARTPool | NM_001080792 | M-019187-01 |
| IER3IP1 | siGENOME Human IER3IP1 SMARTPool | NM_016097 | M-018948-02 |
| TMEM43 | siGENOME Human TMEM43 SMARTPool | NM_024334 | M-014342-01 |
| HECW2 | siGENOME Human HECW2 SMARTPool | NM_020760 | M-007192-00 |
| VPS50 | siGENOME Human VPS50 SMARTPool | NM_024553 | M-012918-00 |
| HTT | siGENOME Human HTT SMARTPool | NM_002111 | M-003737-02 |
| CC2D1A | siGENOME Human CC2D1A SMARTPool | NM_017721 | M-015744-01 |
| MAN1B1 | siGENOME Human MAN1B1 SMARTPool | NM_016219 | M-019670-01 |
| INPP5E | siGENOME Human INPP5E SMARTPool | NM_019892 | M-020852-00 |
| LRRK2 | siGENOME Human LRRK2 SMARTPool | NM_198578 | M-006323-02 |
| RIC1 | siGENOME Human RIC1 SMARTPool | NM_020829 | M-026110-01 |
| VPS13B | siGENOME Human VPS13B SMARTPool | NM_181661 | M-012873-01 |
| REEP1 | siGENOME Human REEP1 SMARTPool | NM_001164732 | M-014235-01 |
| BAP1 | siGENOME Human BAP1 SMARTPool | NM_004656 | M-005791-00 |
| SERAC1 | siGENOME Human SERAC1 SMARTPool | NM_032861 | M-015026-00 |
| CUL3 | siGENOME Human CUL3 SMARTPool | NM_003590 | M-010224-02 |
| STAMBP | siGENOME Human STAMBP SMARTPool | NM_201647 | M-012202-01 |
| LMBRD1 | siGENOME Human LMBRD1 SMARTPool | NM_018368 | M-017956-00 |
| USP7 | siGENOME Human USP7 SMARTPool | NM_003470 | M-006097-01 |
| YIPF5 | siGENOME Human YIPF5 SMARTPool | NM_001024947 | M-018962-02 |
| HACE1 | siGENOME Human HACE1 SMARTPool | NM_020771 | M-007193-01 |
| NECAP1 | siGENOME Human NECAP1 SMARTPool | NM_015509 | M-017872-02 |
| USP7 | siON-TARGETplus Human USP7 SMARTPool | NM_003470 | L-006097-00 |
| USP7 | siON-TARGETplus Human USP7-1 | NM_003470 | J-006097-05 |
| USP7 | siON-TARGETplus Human USP7-2 | NM_003470 | J-006097-06 |
| USP7 | siON-TARGETplus Human USP7-3 | NM_003470 | J-006097-07 |
| USP7 | siON-TARGETplus Human USP7-4 | NM_003470 | J-006097-08 |
| VPS35 | siGENOME Human VPS35 SMARTPool | NM_018206 | M-010894-00 |
| TRIM27 | siGENOME Human TRIM27 SMARTPool | NM_006510 | M-006552-01 |
| RAB4A | siGENOME Human RAB4A SMARTPool | NM_001271998 | M-008539-01 |
| RAB11A | siGENOME Human RAB11A SMARTPool | NM_001206836 | M-004726-02 |
| RAB35A | siGENOME Human RAB35A SMARTPool | NM_001167606 | M-009781-00 |
| SNX27 | siGENOME Human SNX27 SMARTPool | NM_001330723 | M-017346-01 |
